# Design principles of the common Gly-X6-Gly membrane protein building block

**DOI:** 10.1101/2025.06.25.661420

**Authors:** Kiana Golden, Catalina Avarvarei, Charlie T. Anderson, Matthew Holcomb, Weiyi Tang, Xiaoping Dai, Minghao Zhang, Colleen A. Mailie, Brittany B. Sanchez, Jason S. Chen, Stefano Forli, Marco Mravic

## Abstract

Protein behavior in lipid is poorly understood and inadequately represented in current computational models. Design and prediction abilities for bilayer-embedded molecular structures may be improved by characterizing membrane proteins’ most frequent, favored structural features to glean both context-specific and general principles. We used protein design to proactively interrogate the sequence-structure relationship and stabilizing atomic details of two highly prevalent antiparallel transmembrane (TM) motifs with Small-X_6_-Small consensus sequences. A fragment-based data-mining and sequence statistical inference method including cross-evolutionary structure-aligned covariance enabled engineering of *de novo* multi-span TM protein assemblies by successfully encoding Gly-X6-Gly and Ala-X6-Ala building blocks. A highly stable glycine-based design’s X-ray structure hosts Cα-H···O=C H-bonding alongside extensive backbone-directed van der Waals packing, idealizing features of this motif in Nature. Data-driven design navigates sequence space to directly inquiry upon how to encode and stabilize vital membrane protein structural elements, facilitating efficacious construction of lipid-embedded architectures of increasing complexity.

**Significance:** Membrane proteins comprised of α-helices pack together within lipid bilayers, establishing stabilities and architectures that brace function and guide evolution. De novo design was used to clarify the consensus sequences and molecular features encoding of one exceedingly common TM helix packing architecture (∼10% of those in membrane folds). Small-X_6_-Small residue patterns were proven to reliably drive minimal proteins into these antiparallel TM geometries, revealing glycine mainchain hydrogen bonding or via fully apolar interfaces can encode the motif. Optimizing steric packing was the most decisive feature amongst the synthetic proteins. New membrane-specific design methods and model molecules are validated in route to outlining this important sequence-structure relationship broadly impacting the membrane proteome and revealing generalized structure-energetic imperatives governing interactions in lipid.

## Introduction

A few unique helix-helix packing geometry are sufficient to describe about half of membrane protein internal architectures(1–3). This basis set of transmembrane (TM) domain building blocks constitutes folded cores, drives oligomerization interfaces, and anchors dynamic regions and functional sites across organisms, proteomes, and functional families. Although these recursive packing units have been previously identified, most of their sequence-structure relationships remain ambiguous – underutilized in clarifying principles of folding in lipid and membrane protein structure prediction and design. Studies of one such TM building block revolutionized the membrane field: i.e. tight parallel-oriented TM spans mediated by Small-X_3_-Small sequences (Small = Gly, Ser, Ala) including G-X_3_-G(4–6). Successful protein design reflects mastery of that motif’s sequence-structure encoding(7–11). By contrast, packing motifs between antiparallel TM helices are exceedingly common yet poorly defined, having limited experimental evidence clarifying sidechain interaction patterns or consensus sequences that reliably encode these important structures.

One lipid-embedded sequence pattern, Small-X_6_-Small, is becoming increasingly correlated with a packing unit of tight left-handed antiparallel helices. This ubiquitous geometry comprises ∼11% of all 2-helix interaction modes in membrane assemblies and is more common than parallel GAS_right_ motifs (Small-X_3_-Small-type)(1) (12–14). Small-X_6_-Small patterns like G-X_6_-G are conserved in many important protein classes including transporters(15), channels(^16^), lipid enzymes(17), and tetraspanins(18) (**Fig. 1A**), often stabilizing local TM helix bundles within folds that undergo global inter-domain conformational changes. When investigated experimentally, G-X_6_-G motifs within the CD36 receptor superfamily(19) and homo-dimeric multi-drug transporters(12, 20) were functionally essential, mediating necessary TM assembly. Likewise, Small-X_6_-Small sequences in TM peptides can drive their self-assembly in model membranes(13, 21, 22). Small residues repeated every 7 residues (i.e. every other helix turn, ‘*a*’ in ‘*abcdefg*’ heptad) line a continuous helix face and mutually interdigitate to mediate extended antiparallel packing surfaces like leucine zippers, but with much closer mainchains distances(23). At glycines (e.g. G-X_6_-G), backbone atoms can come near enough for potential inter-helix Cα-H···O=C hydrogen bonds (H-bonds), with distinct antiparallel configurations understudied compared to those mediating parallel G-X_3_-G motifs(4, 5). In Nature, however, Small-X_6_-Small sequences do not always utilize H-bonds and in multi-spanning proteins they are seen in diverse contexts including packing both parallel and antiparallel interfaces(24). Thus, it remains uncertain how specific TM geometries (i.e. parallel/antiparallel, oligomer size) are encoded and how polar versus apolar interactions determine energetics for the scope of Small-X_6_-Small sequences.

**Figure 1.**
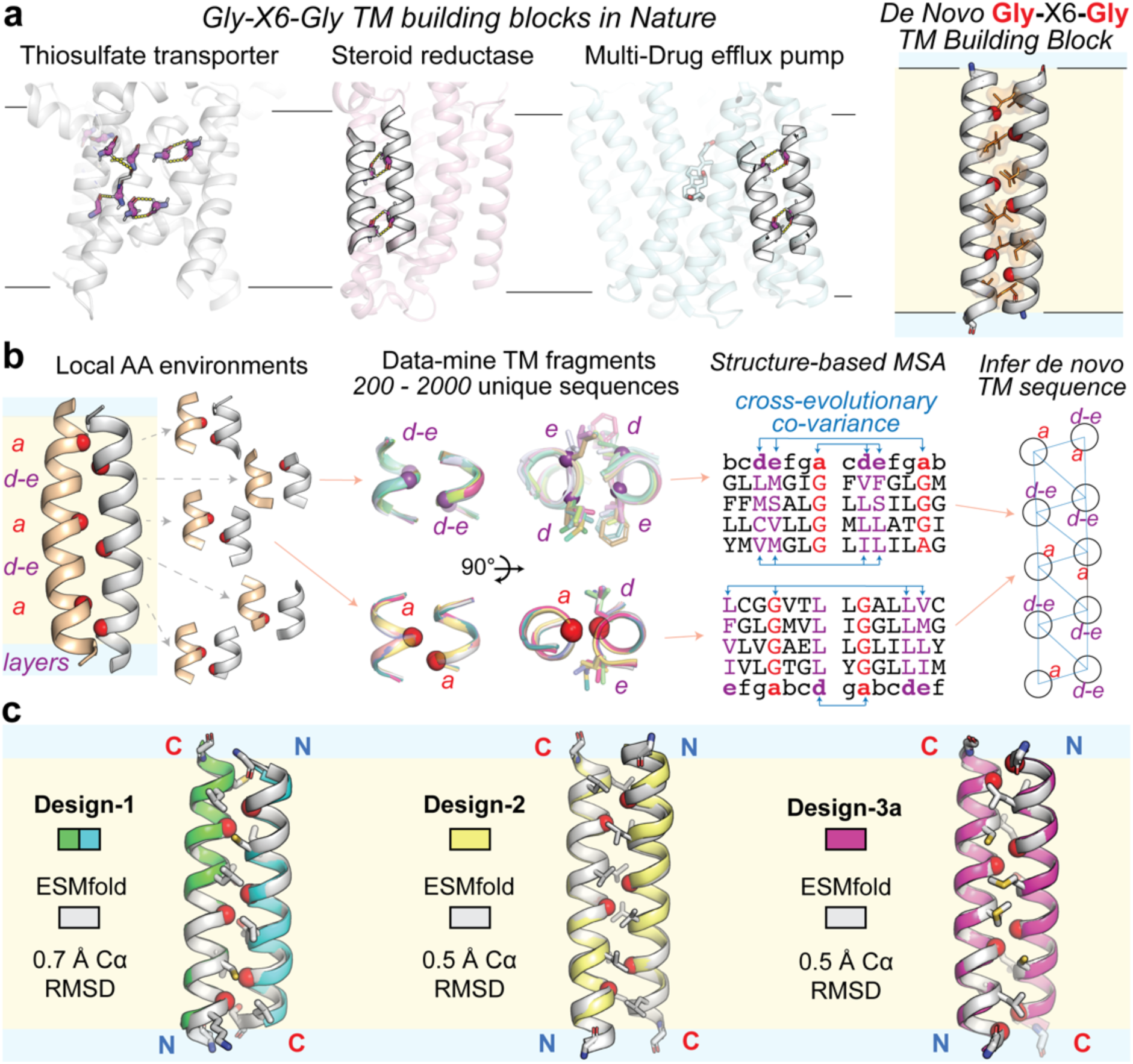
De novo design of minimal TM proteins idealizing the common G-X_6_-G structural building block. (**a**) Motif of close antiparallel TM helices in membrane proteins of the G-X_6_-G repeat (white cartoon); glycines purple sticks; Cα-H hydrogen bonds, yellow dash; PDB codes left to right: 6LEO, 7C83, 4ZP0, 8SRN. Right, design of *de novo* single-span self-assembling dimeric protein representing this TM building block (glycine, red spheres). (**b**) A design model is broken into fragments centered around each interface amino acid (AA), namely ‘a’ and “d-e” packing layers (red and purple Cα spheres, respectively). Natural membrane protein structures (Table S1) are data-mined for instances of similar local helix-helix geometries. The corresponding unique primary sequences found at the matching instances are compiled into a structure-based multiple sequence alignment (MSA) that spans organisms, folds, and functional families across evolution. MSAs from each packing layer are sampled using a graph-based network (right) to statistically infer idealized *de novo* TM protein sequences from natural patterns to find sequences stabilize the local geometries and overall folded structure. (**c**) Computational models of *de novo* antiparallel G-X_6_-G TM building block homodimer complexes overlayed with the ESMfold predicted structures (pLDDT > 0.8). N- and C-termini labeled, blue and red respectively.

Protein design has preliminarily explored the sequence-structure linkage for Small-X_6_- Small patterns(25, 26). Studies of 3 synthetic A-X_6_-A TM peptides find this motif can encode monomers(21), antiparallel dimers(13), or mixed topology trimers(8) – indicating the residues outside the Small ‘*a*’ position heavily influence the fold. S-X_6_-S TM peptides with (SxxLxxx)_3_ repeats assembled primarily to parallel dimers in model membranes(21), mirroring behavior of the mouse erythropoietin receptor (mEpoR)’s similar TM sequence pattern(27). From limited evidence, G-X_6_-G repeats may have greater antiparallel orientation tendancy(13). G-X_6_-G TM peptides mimicking sequences of the conserved 4^th^ TM spans (TM4) mediating subunit dimerization have been used to inhibit small multidrug resistance (SMR) efflux pumps(28). Their activities across SMR orthologs(29) suggest that ‘*d’* and ‘*e’* positions at intermediate helix turns between glycines tune specificity. We previously designed *de novo* TM proteins that selectively recognize and inhibit mEpoR by accommodating mixed Small-X_6_-Small identities (Ala/Ser/Gly at ‘*a*’) and explicitly optimizing ‘*d*’ and ‘*e*’ residues(14). Notably, two recent studies attempting to design parallel TM domain assemblies inadvertently used Small-X_6_-Small patterns; both structures instead adopted corresponding antiparallel TM interfaces(8, 30). Overall, general principles connecting Small-X_6_-Small sequences and antiparallel TM structures are not clear from these limited examples. Filling the major knowledge gaps, i.e. the motif’s structure-energetic basis and impact of non-*‘a’* sequence context in stability and conformational specificity, should broadly impact the understanding of membrane proteins folding and boost capabilities for engineering TM architectures.

Here, we developed a data-driven method we expected would successfully design model TM proteins which encode this antiparallel motif, thus enabling experiments clarifying determinants of folding and stability. The engineered TM sequences hosting G-X_6_-G and A-X_6_-A patterns all fold selectively to the homo-dimeric antiparallel assemblies intended. Characterization of the synthetic G-X_6_-G TM protein family revealed a range of folding free energies and characteristic inter-helical Cα-H···O=C H-bonds. Van der Waals (vdW) sidechain packing at ancillary interacting positions were the dominant molecular feature tuning the specificity and stability amongst the synthetic TM protein variants studied. The algorithm and biophysical principles presented here establish new frameworks to design TM interfaces, clarify membrane proteins principles through the lens of their fundamental building blocks, and construct of more complex synthetic assemblies in lipid by-parts.

## Results

### Computational design of minimal small-X6-small proteins

We sought to design simple single-span TM domains as idealized representative of Small-X_6_-Small TM building blocks in Nature to test and clarify consensus elements encoding their folding. Force field-driven rotamer trials like RosettaMembrane do not recapitulate natural preferences at TM domain interfaces(8, 31). Machine learning models trained on predominantly water-soluble proteins likely undervalue lipid-embedded sequence-structure relationships. Instead, we hypothesized this encoding could be achieved by implementing a very simple framework of statistical inference, inspired by refs (32–34) for globular proteins, and utilizing the relatively limited set of membrane protein structures available. In this fragment-based analysis, we deconstruct the target protein architecture we aim to engineer into smaller subsections of local tertiary geometry, i.e. pairs of mutually packing α-helices (**Fig. 1b**). To determine amino acid identity preferences and inter-residue relationships within each residue’s local environment, we search all known unique membrane proteins for geometrically similar instances of the intended helix-helix arrangement (1 Å backbone RMSD, **Table S1**) and collect all primary sequences observed to support the conformation. This results in structure-based multiple sequence alignments (MSA) of interacting groups surrounding each key interface position (**Fig. 1b**). We posited that a simple 2 term scoring function comparing observed per-residue MSA statistics to expected frequencies would be sufficient to discern amino acid and interaction group choices that can stabilize the intended fold: amino acid enrichment (1-body) and co-variance between inter-helical residues (2-body). To search sequences globally optimizing this statistical score, a graph-based representation of residues constituting the interface was used (**Fig. 1b**), thus generating full *de novo* TM domains with native-like molecular features and patterns likely for the structure.

Our generative design model aims to similarly leverage co-variance’s predictive power proven in emerging AI/ML models(35–37) by mining similar nano-environments from unrelated membrane proteins across diverse organisms, protein families, and topologies. The approach overcomes the lack of relevant protein familial MSA data crucial to those models, done by constructing analogous structurally aligned “cross-evolutionary” MSA data and identifying convergent molecular patterns for *de novo* design. Thus, we hypothesize that the quantity of existing membrane-embedded structures combined with the proposed data structure (sequence-structure tertiary fragment pairs) are sufficient to capture and proactive engineer TM-specific relationships – presumably with interaction strengths comparable to or optimized beyond those in Nature. Designing new minimalist TM proteins serves as a simple accessible model experimental system. Each sequence is a data-driven hypotheses of features suitable for that protein topology. We explored this design-test-learn platform to evaluate the computational approach and probe consensus relationships between sequence, structure, and energetics for TM building blocks like Small-X_6_-Small.

We benchmarked the new algorithm for encoding sequences compatible with our recently designed membrane pentamer channel, which hosts a common parallel-oriented building block(38). We previously showed 1-body per-residue amino acid enrichment from this cross-evolutionary structure-based MSA analysis has predictive power of highly stable protein variants. Our new generative model now including a 2-body residue co-variance term results in superior recovery of sequences known to have experimentally proven stable folding, notably improving correlated pairwise amino acid choices at residues with larger Cα-Cα distances (**Fig. S1**).

Next, we deployed this algorithm to design 1-pass TM spans hosting Small-X_6_-Small repeats testing if resulting sequences can stabilize homo-dimeric folding to the tight packing antiparallel motif(3). First, the range of left-handed backbone arrangements relevant to this motif in Nature was defined by sampling a set of idealized coiled-coiled molecular models varying in helix-helix geometry(39) and scoring each by frequency of occurrence in known membrane proteins(40) (**Fig. 1b, S2; Table S1**). We selected 4 common, but mutually distinct representatives to design: 3 close backbones (8.0 or 8.4 Å inter-helical distance) preferring G-X_6_-G patterns and a wider backbone (8.8 Å) preferring A-X_6_-A patterns **(Fig. 1c, S2b**).

For protein design, a set of sequences statistically optimized for each inter-helix geometry was produced (logos in **Fig. S3**). Then, top ranked sequences were subsequently modelled and repacked in all-atom detail. Models with inevitable clashes were discarded. Designed sequences with <2 Å RMSD and RMSF stability all-atom lipid bilayers molecular dynamics (MD) simulations were selected for experimental characterization. We additionally generated a structure-based manually designed sequence, G-X_6_-G Design-1, using the same starting backbone used for the statistically optimized G-X_6_-G Design-2. This compares our computational approach versus rationale design for efficacy to derive G-X_6_-G sequences that fold, the key differences being “*d*” and “*e*” sidechains (**Fig. S2b**). Five unique parent *de novo* TM proteins resulted, varying in hydrophobic length, composition, and number of repeats (**Fig 2a**, **Table S2**).

**Figure 2.**
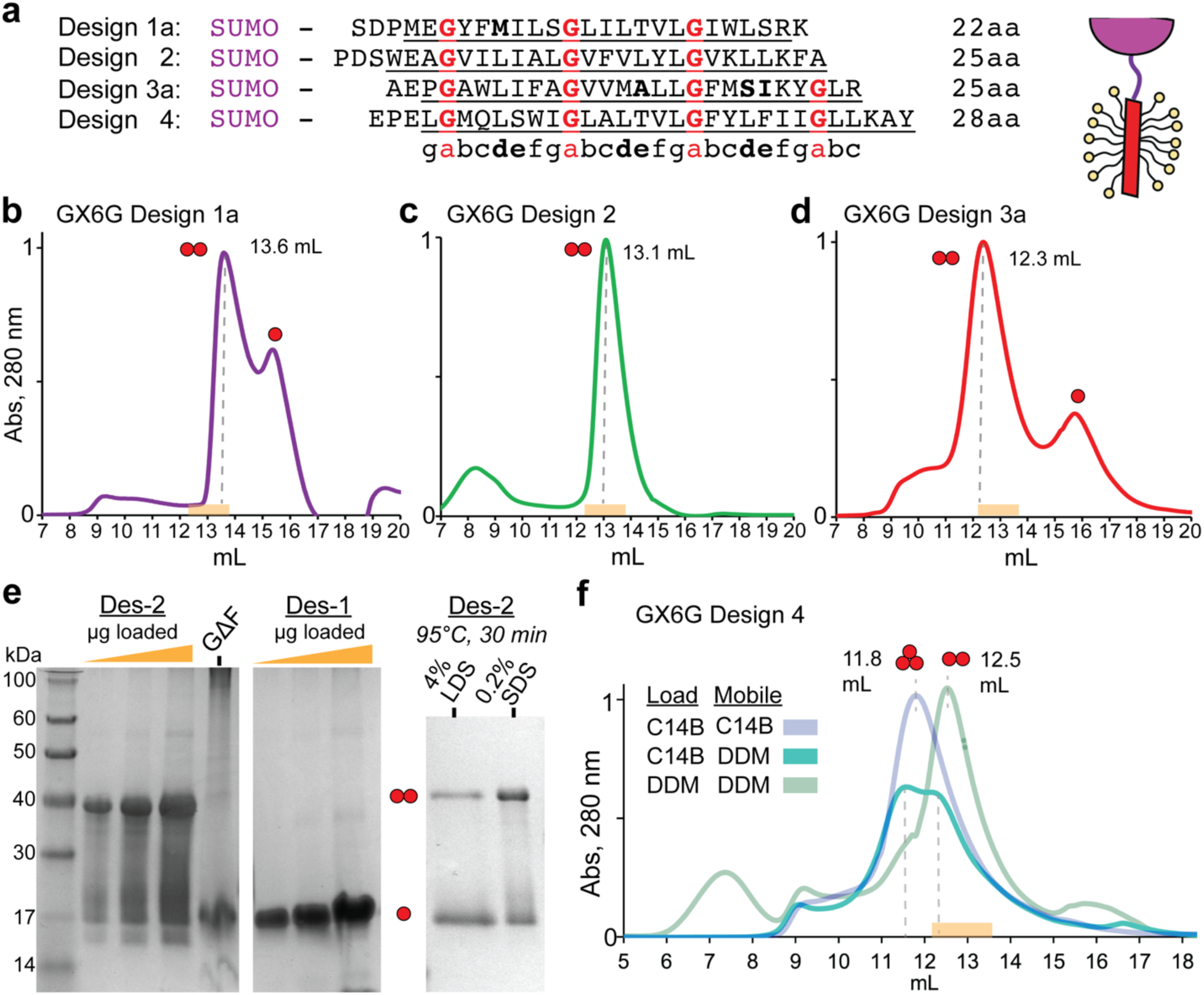
Folding of synthetic G-X_6_-G TM domains as SUMO fusion proteins. (**a**) N-terminal SUMO fusion constructs for synthetic G-X_6_-G TM domains aligned by heptad position. Glycines red, bolded. Interface positions, bold. TM domain length listed right. Left, cartoon of SUMO (purple) and TM domain (red). (**b**) Size exclusion chromatography (SEC) trace of absorbance 280 nm (normalized to peak max) by column volume of 2 mg/mL SUMO-TM Design-2 (representative of n=3) on superdex200i 10/300 column, mobile phase: 50 mM Na Phosphate pH 7.8, 100 mM NaCl, 0.5 mM EDTA, 1 mM DTT, and 1 mM DDM. Red spheres, assigned oligomer. Orange box is expected elution range for a dimer. (**c**) SEC of SUMO-Design-2 (representative of n=3) in 1 mM DDM, major peak falls into dimer range (**d**) SEC of SUMO-Design-3a (representative of n=3) in 1 mM DDM, major peak falls into dimer range (**e**) Left, SDS-PAGE of 1 mg/mL Design-2 in DDM at increasing loading volumes in Tris-Glycine-SDS or the point mutant with central Gly mutated to Phe (G16ΔF). Middle, Design-1a gel. Right, Design-2 after mixing with 4% w/v lithium dodecyl sulfate (LDS) or 0.2% SDS and heating to 95 ° C for 30 minutes. (**f**) SEC of SUMO-Design-4 incubated in 30 mM C14B or 40 mM DDM and run in mobile phase of 3 mM C14B or 1 mM DDM. C14B-incubated protein in C14B mobile phase results in a likely trimer (11.8 mL). DDM-incubated protein in DDM mobile phase results in a dimer. C14B-incubated protein in DDM mobile phase results in trimer-dimer mixture.

Additional variants were derived from the guiding designs models to facilitate cysteine-directed folding experiments and to test contributions from *‘d’* and *‘e’* interface residues. G-X_6_-G Design-1 was produced as the original cysteine-containing design (variant 1b) and as a methionine point mutant (variant 1a, C11M), expecting analogous sidechain packing at this *‘d’* residue. Three variants of G-X_6_-G Design-3 were designed to assess contributions of potential inter-helical H-bonds. The original Design-3 (i.e. variant 3a) hosts Ser19 at ‘*d*’, expecting an inter-helical H-bond to Leu5’s backbone carbonyl (**Fig. S4a**). Design-3b varies from 3a by the synonymous Ser19Cys mutation, retaining that intended inter-molecular H-bond, and mutating Ala13Ser at ‘*e*’ to add another inter-molecular H-bond. A more drastically mutated variant 3c further challenged the structure’s tolerance, replacing 3a/b’s Ser/Cys19 H-bonding at ‘*d*’ with Met19’s apolar packing and adding back a putative inter-helical H-bond at the preceding ‘*d*’ Met12Ser (**Fig. 4b**). Conversely, these Ser/Cys sidechains could prefer intra-helical rather than inter-helical H-bonds.

The final stable design models for the G-X_6_-G and A-X_6_-A sequences were compared with the range of antiparallel TM geometries occurring naturally and with those predicted *ab initio* by orthogonal methods. A previous clustering analysis of all ways 2 helices interact in membrane proteins identified the tight left-handed antiparallel motif we designed as the 3^rd^ most common recurring packing geometry (∼141 unique helix-helix pairs)(1). The 5 synthetic TM proteins’ expected backbone conformations adopt native-like variations of local fold family and should thus appropriately represent its sequence-structural relationship (**Table S3**). Predicted structures by AlphaFold-3 (AF3)(37) were confident but likely incorrect for 4/5 TM sequences, adopting low contact density parallel-oriented helices (**Figure S5**). AF3 did accurately predict G-X_6_-G Design-4 (2 Å RMSD), which has the longest TM span. ESMfold(35) predicted high-confidence models with <0.7 Å backbone RMSD agreement for each design, suggesting the intended fold is the lowest energy structure for each sequence and this large language model (LLM) recognizes membrane-spanning Small-X_6_-Small motifs (**Fig. 1c**, **Fig. S4b**).

### Stable and specific folding of de novo small-X_6_-small TM proteins

The ability of the designed TM domains to drive stable specific folding to the intended homo-dimer assembly was assessed by size exclusion chromatography (SEC), using SUMO fusion proteins of the G-X_6_-G variants and characterizing each design’s oligomeric distribution in multiple detergents: n-Dodecyl-B-D-Maltoside (DDM), Myristyl sulfobetaine (C14B), and N,N-dimethyldodecylamine N-oxide (LDAO). A set of TM domains known to dimerize(13, 41) and trimerize fused to SUMO established reference SEC migration and folding behavior, eluting at expected detergent-oligomer complex sizes in different micelles and achieving adequate resolution (≥0.5 mL separation) to assign states for the designed TM oligomers (**Fig. S6b-c, Table S2**). Design-1a and −1b migrate as a 2-species mixture in both DDM and LDAO, assigned as dimer and monomer peaks based on size, with apparent ∼60-75% fraction folded dimer – reflecting a modest stability and equivalency for Cys11 or Met11 at *‘d’* (**Fig. 2b, S6a**). By contrast, Design-2 robustly folded exclusively to a dimer in DDM (>98%, no detectable monomer, **Fig. 2c**). Design-2, −3a and −3b were predominantly dimeric (>80%) in all detergents tested (**Fig. 2c-d; S6d-f**). Design-3c variant exhibited much lower folding specificity, eluting as a mixture of dimer and a larger likely trimeric oligomer: indicating G-X_6_-G does not strictly encode TM dimerization (**Fig. S6f**). Thus, the 3 parent G-X_6_-G TM designs encode specific homo-oligomeric folding to the dimeric state, exhibit tolerance and disruption as the intended interface is varied, and have distinct monomer-dimer equilibria.

Next, relative stabilities were assessed. After exposure to harsh conditions expected to denature most natural membrane proteins, Design-2 remained predominantly dimeric in SEC indicating high stability: incubation with excess short-chain detergent (200 mM octyl-glucoside, **Fig. S6d**) or heating at 85-90° C in LDAO or DDM (**Fig. S6d**, **Fig. S7a**). Design-3a exhibited similar amounts of folded dimer after 15 minutes at 85° C in LDAO (**Fig S6e**). The Design-2 TM protein shows greater resistance to denaturation, running as a sodium dodecyl sulfate (SDS)-resistant dimer by poly-acrylamide gel electrophoresis (PAGE), whereas Designs-1 and Design-3 variants run as monomers (**Fig 2e**, **Fig. S7b**). Mutation of the central motif glycine (G16F) ablates this dimerization, indicating responsibility of the G-X_6_-G sequence in its assembled structure. Incubation at high concentration of lithium dodecyl sulfate (LDS) partially reduced dimerization in gel migration, showing folding’s detergent-dependence. Heating in SDS had minimal impact to Design-2’s dimeric folded fraction by PAGE (**Fig 2e**), indicating either temperature-independence or rapid dimer refolding. Thus, Design-2 is exceptionally resistant to denaturation, amongst the most stable membrane proteins characterized to date.

Design-4, which has the longest hydrophobic span (4 G-X_6_-G repeats), showed strong self-assembly like the other designs, but differed as its oligomeric state was detergent-dependent (i.e. folding specificity). When incubated in excess DDM and ran in DDM mobile phase, Design-4 instead folds to a dimer as intended (12.8 mL, **Fig. 2f**). When purified in C14B and ran in SEC with DDM mobile phase, Design-4 eluted as two peaks – a likely dimer and a slightly larger species. Running C14B-purified protein in C14B mobile phase yielded a monodisperse peak of this slightly larger species (11.8 mL). Glutaraldehyde cross-linking confirms a trimer dominates in C14B while a dimer is preferred in DDM (**Fig. S8**). To discern the molecular basis, we predicted the trimer’s structure. ESMfold’s top model bears only a tight antiparallel dimer and a fully dissociated 3^rd^ TM span (i.e. non-trimer) whereas AF3 predictions are low confidence – neither clarifying the alternative stable trimer conformation (**Fig. S2a**). While Design-4’s dimerization is achieved in DDM, off-target folding in C14B highlights that G-X_6_-G motifs can stabilize specific trimers as well as dimers depending on the sequence and model membrane context. We interpret this detergent-dependent behavior observed as the interplay between interaction energetics and solvation, i.e. TM hydrophobic thickness matching.

We next addressed whether Designs 1-3 encode the antiparallel orientation using thiol-disulfide equilibrium exchange(42). N- and C-terminal cysteine-tagged TM peptides of each design were mixed in a 1:1 ratio (**Fig. 3**, **Table S2**), reconstituted in aqueous glutathione redox buffer with 100 molar excess C14B (>1 micelle per peptide) for overnight reversible oxidation, then read out by LC-MS. We expected formation of the disulfide-bonded homo- or hetero-dimers to reflect the underlying preference of noncovalent TM interaction geometries, parallel versus antiparallel (**Fig. 3a**). For Design-1a, −2, and −3b, the dominant disulfide-bonded dimer species is consistently the antiparallel hetero-dimer (**Fig. 3b-c**, **Fig S9-11**), although monomer fraction varied between designs. The two possible covalent parallel homo-dimers were either undetectable or present at much lower abundance (at least 5-fold less). These *de novo* G-X_6_-G TM domains all demonstrate strong non-random preference for the antiparallel geometry, matching the topology intended by design.

**Figure 3.**
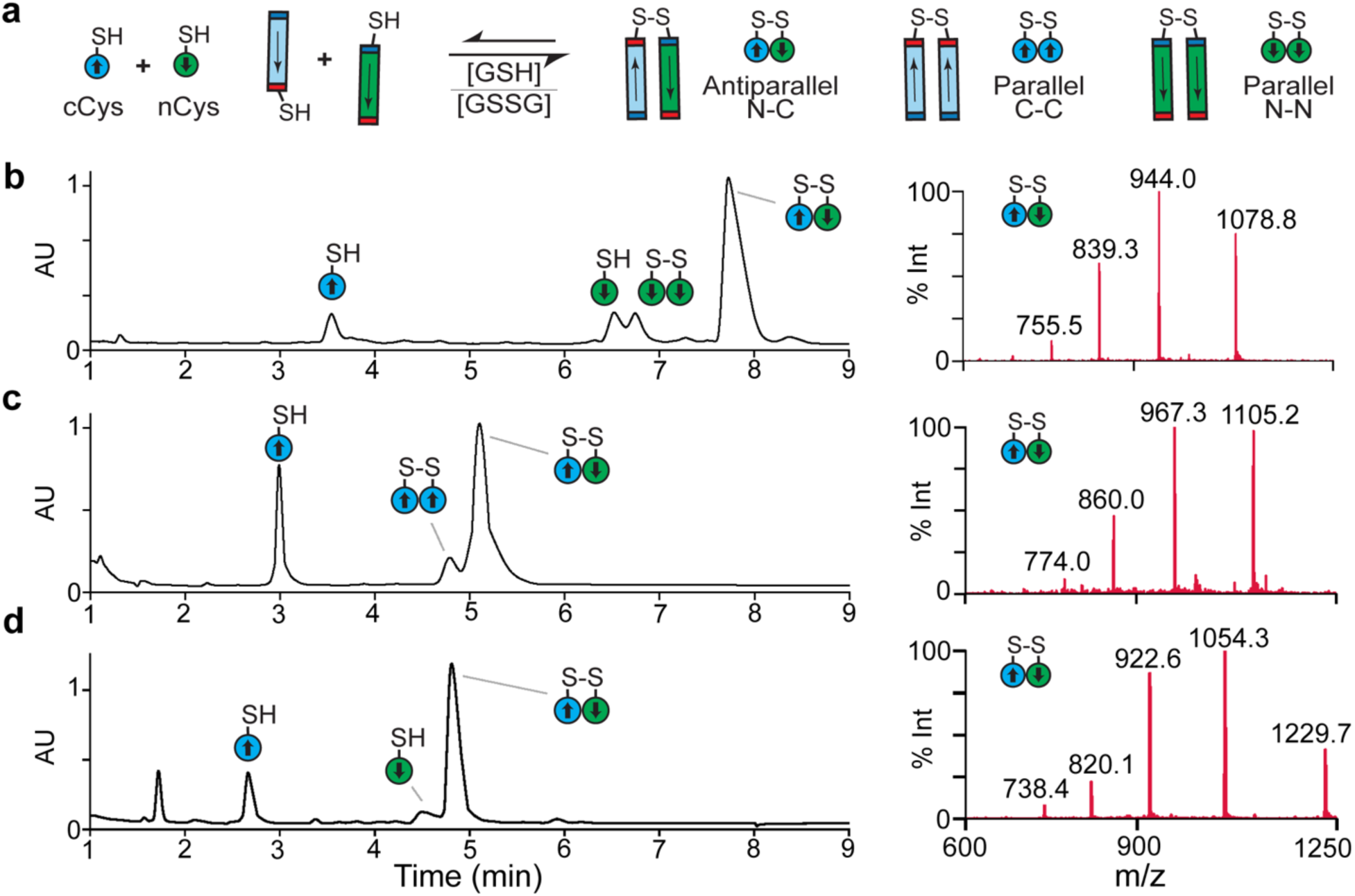
Antiparallel interaction topology of G-X_6_-G TM domains by thiol-disulfide equilibrium exchange. **(a)** N-terminal and C-terminal cysteine (nCys, cyan; cCys, green) TM peptides are reconstituted at 50 µM concentration in aqueous 1.5 mM glutathione buffer (3:1 ratio [GSH]:[GSSG]) with 10 mM C14B results in equilibrium oxidation into 3 possible disulfide species: parallel cCys homodimer, parallel nCys homodimer, and antiparallel N-C heterodimer. **(b)** UV max plot reverse-phase high-performance liquid chromatography (RP-HPLC) trace of Design-1a reaction (representative of n=3), left. TM peptide monomer and covalent dimer species denoted by cartoon. Right, mass spectra of the major antiparallel N-C heterodimer species (expected: 7544.8 Da, Observed: 7544.2 Da). **(c)** UV trace (left) and mass spectra (right) from LC-MS of Design-2 thiol-disulfide equilibrium exchange reaction showing the antiparallel N-C heterodimer is the major disulfide bonded species (expected: 7729.2 Da, Observed: 7729.1 Da). Representative of n=3. **(d)** LC-MS of Design-3a thiol-disulfide equilibrium exchange reaction and major antiparallel N-C heterodimer (expected: 7372.8 Da, Observed: 7372.8 Da) Representative of n=3.

**Figure 4.**
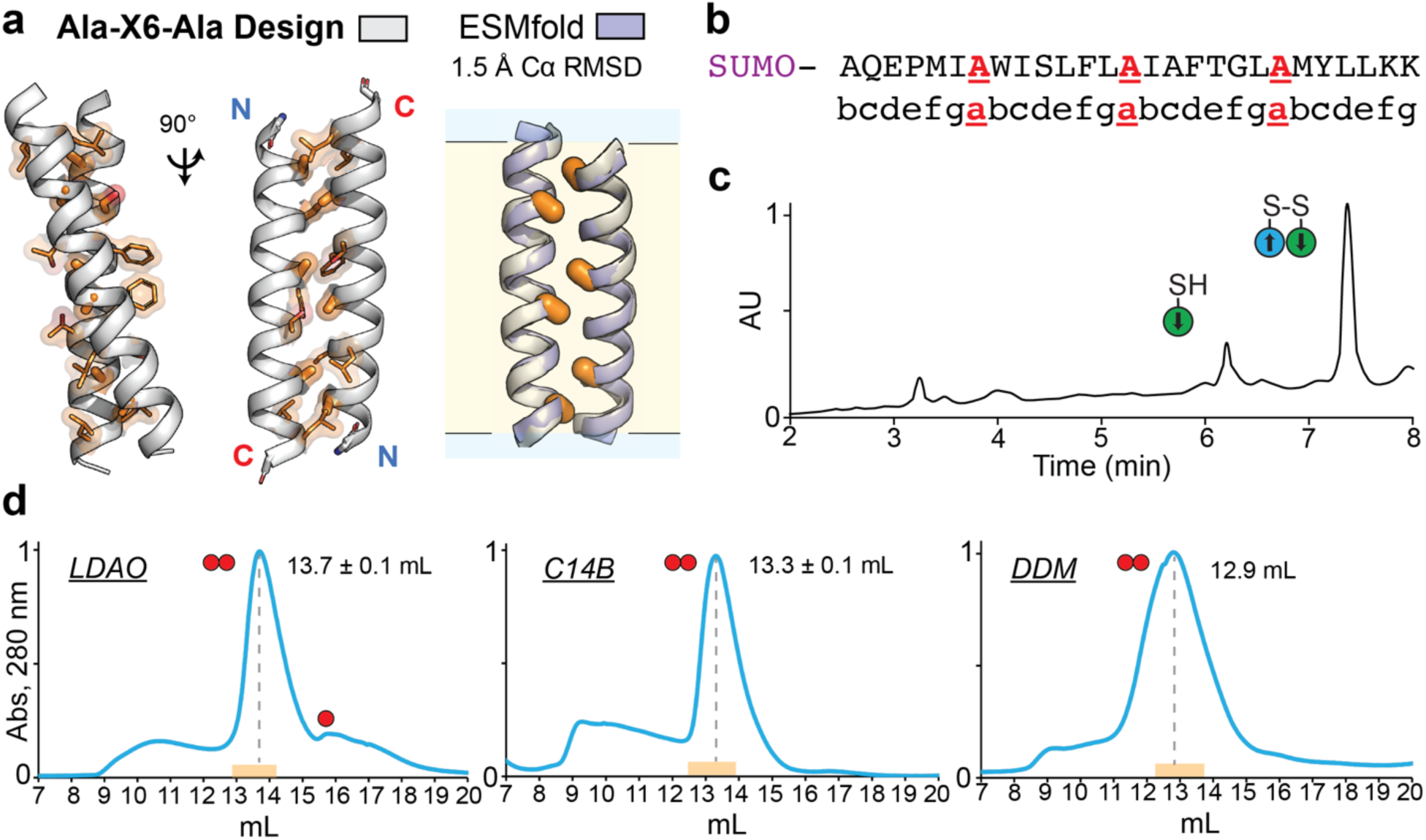
Folding of synthetic A-X_6_-A TM domain protein. (**a**) Design model of *de novo* A-X_6_-A TM domain forming antiparallel fold (white), and overlayed with ESMfold structure prediction (purple, 1.5 Å Cα RMSD). Motif alanines, orange sticks. (**b**) Designed A-X6-A sequence. Motif Ala residues underlined in red, aligned with 7-residue heptad repeat. (**c**) Thiol-disulfide equilibrium exchange LC-MS UV maxplot chromatogram of A-X_6_-A nCys and cCys peptide mixture reconstituted in 10 mM LDAO, with major peaks nCys monomer and N-C heterodimer annotated by cartoon as in Figure 3, representative of n=3 trials. (**d**) Size exclusion chromatography (SEC) trace of absorbance 280 nm (normalized to peak max) by column volume of 2 mg/mL SUMO-TM A-X_6_-A design (representative of n=3) on superdex200i 10/300 column, mobile phase: 50 mM Na Phosphate pH 7.8, 100 mM NaCl, 0.5 mM EDTA, 1 mM DTT, and detergent (6 mM LDAO, 3 mM C14B, and 1 mM DDM). Red spheres denote monomer or dimer assignment of peak. Orange box is expected elution range for a dimer in that detergent based on known TM domain reference proteins.

### A-X_6_-A Design forms antiparallel dimers

Unable to form Cα-H···O=C H-bonds due to stereochemistry and distance, A-X_6_-A motifs are expected to achieve slightly wider variations of this TM building block geometry analogous to water-soluble “Ala-coils”(43, 44). Thus, our *de novo* A-X_6_-A Design predominantly utilizes apolar sidechains with tight geometrically compatible vdW interactions; S12 at *‘d’* forms intrahelical H-bonds, packing its Cβ methylene. Motifs alanine methyl “knobs” project deeply into the vacant backbone polar “hole” region between inter-helical ‘*d*’ and ‘*e*’ residues (**Fig 4a-b**). The ESMfold prediction adopts identical sidechain packing and inter-helix distance (∼8.8 Å) as the design model (1.5 Å RMSD). Packing small residues in TM spans may also result in entropy from lipid exclusion from the more exposed polar mainchain upon folding, leading to favorable solvo-phobic effects – reciprocal to the hydrophobic effect in water(26, 45, 46) – although its energetic significance and magnitude for different small residues (G, A, S) is unclear.

The SUMO-fused A-X_6_-A TM domain drives specific folding to a monodisperse dimer, which is not SDS-resistant in gel migration, thus weaker than G-X_6_-G Design-2 and similar to Design-3 (**Fig 4d**, **Fig. S7b**). The minimal A-X_6_-A as well as the G-X_6_-G TM peptides reconstituted in mild detergent form SDS-resistant oligomers (>6 kDa) during SDS or lauryl sarkosyl PAGE(47) (**Fig. S4**). Although assignment of oligomeric states was difficult due to anomalously migration(48), the TM domains readily self-assemble robust to SDS. Thiol disulfide exchange of N- and C-terminal A-X_6_-A Design TM peptides reconstituted in LDAO or C14B reveals the antiparallel heterodimer as the predominant disulfide-bonded dimer species, indicative of strong antiparallel preference (**Fig. 4c**, **Fig S13**). Thus, our generative model implicitly optimized steric packing of apolar character to encode the wider backbone variant of the TM geometry accommodating A-X_6_-A, lacking the glycine’s close approach or H-bonds. Likewise, our design’s ‘*d*’/‘*e*’ sidechains achieve structural specificity to antiparallel dimerization, differentiated it from past synthetic A-X_6_-A sequences which adopt multiple or off-target folds(8, 21).

### Structural basis of Design 2’s high stability

To discern the basis of Design-2’s stability, we solved a 3.27 Å X-ray structure in octyl-glucoside (**Fig. 5, Table S4**). All 3 dimers in asymmetric unit have sub-atomic accuracy to the design model (0.6±0.1 Å RMSD, **Fig. S14**). As expected, sequential G-X_6_-G motifs mutually line an extended antiparallel interface of left-handed super-coiling α-helices, contrasting G-X_3_-G motifs’ shorter X-crossed interfaces(5, 41) (**Fig. 5a**). Tighter than expected inter-helical distances (7.5±0.2 Å) yield close inter-glycine distances and favorable backbone-backbone vdW surfaces. Glycine Cα-H_2_’s act as non-canonical “knobs” packing into the backbone “hole” comprised of inter-helical residues ‘*d*’, ‘*e*’, and the subsequent ‘*a*’ glycine (*i+3*, *i+4, i,* respectively). Design-2’s apolar ‘*d*’-‘*e*’ sidechains pack with apparent mainchain-directed “knobs-to-holes” interactions, achieved similarly for Leu or beta-branched Ile or Val, burying the expanded polar backbone area flanking each motif glycine (‘*g*-*a*’ and ‘*b*-*a*’ “holes” with *i*-1 and *i*+1 neighbors, respectively) (**Fig. 5a**). Glycines repeating on the same helix face in the unfolded state TM monomer exposes large polar mainchain surfaces (**Fig. 5b**) likely putting strain on nearby apolar lipid tails which, upon glycine burial during folding, should be released to the lipid bulk regaining favorable entropy. Design-2’s heightened stability appears correlated with its exclusive use of large sterically compatible apolar ‘*d*’-‘*e*’ amino acids: certainty due to extensive enthalpic vdW surfaces from intimately packing backbone grooves and possibly entropically from sealing ample backbone polar area from lipid. Designs-1 and −3 utilize more small or polar sidechains (Ala, Ser, Cys, Thr), with some intended for inter-helix H-bonding, but ultimately resulted in weaker apparent folding.

**Figure 5.**
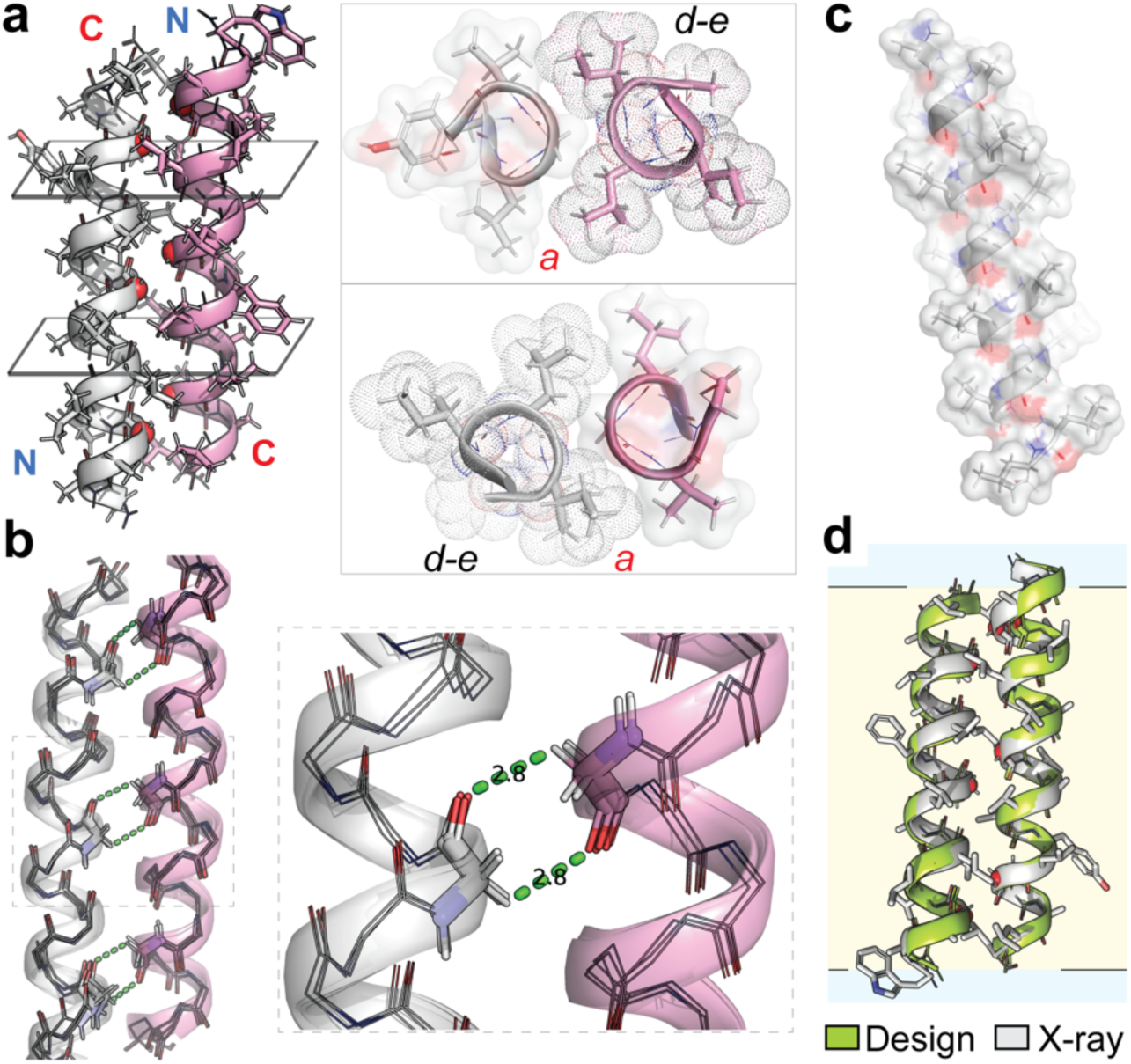
X-ray crystal structure of Design-2 G-X_6_-G TM antiparallel dimer complex. (**a**) Atomic structure of G-X_6_-G protein Design-2 antiparallel complex (N- and C-termini denoted) packing tightly at glycine residues (red spheres). Glycine residues at ‘a’ positions form “knob-to-holes” packing into the backbone hole between apolar ‘d’ and ‘e’ residues. Sidechains of ‘d’ and ‘e’ adopt backbone-directed packing into backbone holes surrounding glycine ‘a’ residues, (knobs to g-a, a-b holes). (**b**) Interhelical Cα H-bonding of glycine protons to carbonyl of symmetric glycine residues, overlaying the 3 unique TM dimer units in the asymmetric unit. (**c**) Extensive exposure of polar backbone atoms of the unfolded state TM domain along a continuous helix face lined helix by glycine7-residue repeats lacking sidechains. (**d**) Structural overlay of Design-2 crystal structure (white) and intended *in silico* design model (lime) of 0.6 Å Cα RMSD.

Design-2 forms clear Cα-H···O=C H-bonds repeating between each motif glycine and the carbonyl of the opposing symmetrically related glycine across the interface, 6 possible per dimer (**Fig. 5c**). Cα-H···O distances are particularly close, 2.6-2.9 Å in the core with a mean distance of 3.0 Å across the 3 unique dimers, including longer distances at termini. These closer H-bonds likely contribute more energy than the common 3.2-3.5 Å Cα H-bonds in TM regions, and are similar to those in glycophorin A’s (GpA) prototypical G-X_3_-G motif (pdb: 5EH4, 3.0**±**0.3 Å, **Fig. S15a**)(49). Designs 1-4 models from Rosetta(50), Charmm36(51), or ESMfold(35) consistently form serial Cα-H···O=C H-bonds, however at further distances (3.1-3.5 Å) indicating these systems undervalue the interaction and are unsuitable for precisely estimating design-specific differences. If each repeating Cα-H H-bond contributes only modestly, up to the −0.9 kcal/mol estimated for a G-X_3_-G motif(52), the cumulative effect should provide substantial folding free energy – as in Design-2.

### Cα H-bonding in natural and synthetic glycine TM building blocks

We next analyzed the characteristics of the glycine Cα-H···O=C H-bonds to discern energetic determinants, particularly Cα-H_2_ proton pseudo-chirality. In Design-2’s three crystallographic dimers (17 H-bonds), one Cα-H adopts a more favorable geometry (2.6-3.0 Å Cα-H···O distance; Cα–H–O angle, ∼110°; H–O–C angle, ∼130°) while the other proton is further but in plausible H-bonding distance at less ideal angles (3.0-3.4 Å Cα-H···O; Cα–H–O angle, 90-100°; H–O–C angle, 90-100°). The Cα-H with stereochemistry analogous to a sidechain (Cα-Cβ) we call “R” (unique to Gly); its pseudo-chiral counterpart the “S” proton (**Fig. 6a**). The “R” proton is the favored H-bond donor in 16 of 17 instances (**Fig. 6b**). This small difference in “R” vs “S” Cα-H···O distance (D_RvS_ = 0.3 **±** 0.1 Å) suggests a possibly geometric balancing to align both protons into carbonyl orbital interactions, influencing these atoms’ positions. This antiparallel H-bonding starkly contrasts the more imbalanced pseudo-chirality in GpA’s parallel G-X_3_-G motif wherein “R” protons are the exclusive H-bond donors (“S” protons, 3.8 **±** 0.4 Å; mean D_RvS_ = 0.8 **±** 0.2 Å, n=8; **Fig S15a**).

**Figure 6.**
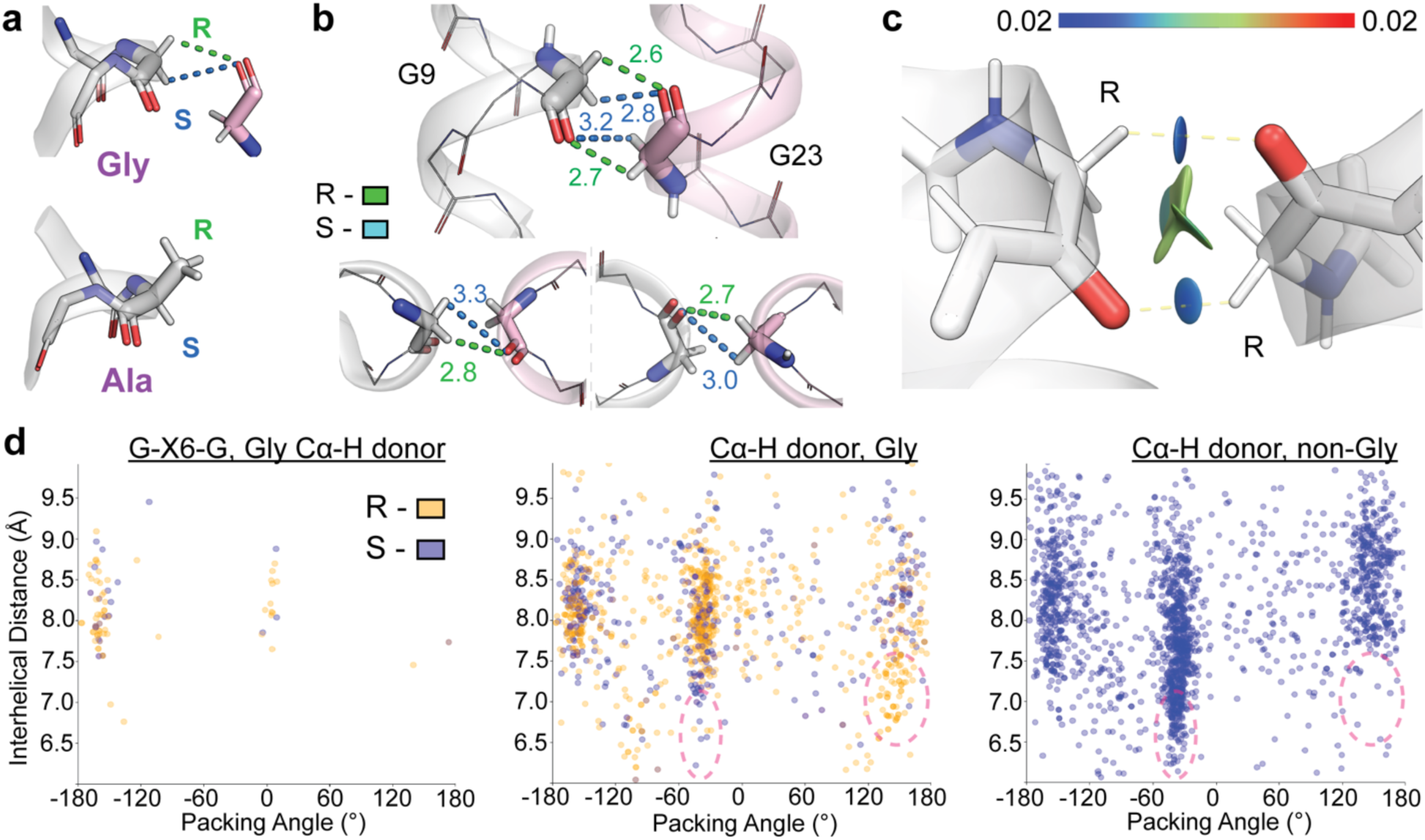
Cα H-bonding characteristics in *de novo* Design-2 and natural membrane protein TM domains. (**a**) Pseudo chirality of glycine protons in 3D environment of G-X_6_-G motif with inter-helical Cα H-bonds. “R” and “S” protons based on the analogy to “L” amino acids sidechain stereochemistry (green and blue, respectively). (**b**) Geometry of R and S glycine protons in interhelix H-bonds in Design-2 crystal structure. Lower, possible bivalent interactions of R and S protons to the inter-helical glycine carbonyl. (**c**) QM optimized structure of close contact glycines in Design-2. Atoms shown as stick; 0.5 a.u. isosurface of reduced density gradient (*s*) is colored by the sign(λ_2_)ρ (a.u.) (color bar), a measure of interaction strength. R protons labelled; partially covalent H-bonds, yellow dotted line. (**d**) Cα H-bonds in natural membrane proteins from a non-redundant structural database (Table S1) characterized by yellow or blue dots for R or S pseudo-chirality and plotted by the local TM domain helix-helix geometry hosting the interacting. Left, Cα H-bond donors in TM domain G-X_6_-G motifs to any acceptor TM domain sequence. Middle, all Gly Cα H-bond donors. Right, all non-glycine Cα H-bond donors. Dotted ovals denote key population differences in middle and left plots.

Next, to investigate “R” and “S” proton H-bonding character, we performed quantum mechanics (QM) calculations on the central Cα-Cα region. Based on the QM minimized structure with harmonic backbone restraints, there is an extensive attractive interface between glycines according to the non-covalent interaction (NCI) index(53) (**Fig. 6c**). Inter-helical Cα “S” proton contributions are largely disperse vdW forces; its electrostatic contribution is minimal, unlikely to impact to protein geometry. Both “R” protons are associated with more localized strongly attracting interactions to inter-helical carbonyl partners having associated bond critical points with properties indicative of H-bonds with covalent character and weakly shared electron density(54), analyzed through Quantum Theory of Atoms in Molecules (QTAIM) framework(55). Based on a published correlation between bond critical point electron density and hydrogen bond strength(56), they are estimated to provide 2.4 to 2.9 kcal/mol per H-bond in stabilization energy *in vacuo* (**Fig S15b-d**). Thus, the balanced Cα-H_2_ arrangement is a consequence of tight mainchain packing vdW surfaces simultaneously positioning “R” protons for near-optimal Cα-H···O H-bond geometries without compromise.

We examined whether Design-2’s Cα-H···O=C features approximate those in natural G-X_6_-G motifs and TM helix geometries (**Dataset S1**). Parallel-right (Small-X_3_-Small), Antiparallel-left (Small-X_6_-Small), and Antiparallel-right (consensus underdetermined) recursively host Cα H-bonds (**Fig. 6d**). Parallel left-handed packing is common(1), but seldomly positions Cα H-bonds.

TM G-X_6_-G sequences predominantly host Cα H-bonds in antiparallel left-handed geometries (−135-165° crossing angle) with 70% (66/94) having balanced “R” proton-led H-bonds matching the pseudo-chirality of Design-2 (**Figs. 6d**). A subset of G-X_6_-G Cα H-bonds occur in near-completely parallel TM helices (crossing angle, ∼0 **±** 15°). However, 95% (20/21) come from a single protein family in highly kinked conformations (5 ATP synthetases in our database with divergent TM spans, <40% global identity) – distinct from ideal coiling helices in Antiparallel-left structures.

Cα H-bonds are ubiquitous across the membrane proteome, with or without glycine residues. Parallel-right geometries (−25-50° crossing angles) favor imbalanced “R” Cα H-bonds with glycines, and closer inter-helix packing (6.5-7.5 Å) can occur at non-Gly amino acids (“S” protons). The latter tight inter-helix “S” proton H-bonds are typically found preceding a glycine at the i-3 position (<u>X</u>xxG; sometimes within a Small-X_3_-Small motif, e.g. Small-<u>X</u>xx-G), proceeding a packing glycine at the i+4 position (G-xxx-X), or when packed into an inter-helical Small-X_3_-Small motif with 1 or 2 glycine. Antiparallel-right topologies (135-165° crossing angle) access closer inter-helical distances when hosting glycine “R” Cα H-bonds (6.5-7.7 Å) compared to non-glycine “S” H-bonded interfaces. Different interhelical geometries in membrane proteins are predisposed for distinct Cα H-bond characteristics. Likewise, natural Antiparallel-left TM building blocks prefer similar balanced-type “R” −directed geometries as in our *de novo* designs.

## Discussion

Here, we benchmark a data-driven design approach and refine the sequence-structure relationship for one of the most common ways membrane proteins pack. In addition to corroborating that G-X_6_-G and A-X_6_-A are consensus patterns able to drive antiparallel TM geometries, our work illuminates novel determinants for encoding this architecture. One defining feature is *‘d’*-*‘e’* sidechains packing directed towards the mainchain with high steric specificity within the tight local inter-helix geometry. Given that diverse sequence compositions and sidechain sterics are compatible at intervening helix-turns amongst designs, evaluating this 3D packing quality within its structural context may be more predictive than simplified sequence-based rules – differing from soluble and TM coiled-coils with strict, clear patterns (e.g. beta-branching)(38, 57, 58). Notably, AI models show discrepancies for this helix-helix geometry, which is seldom found in water-soluble folds(1). For evaluating apolar TM packing structures in membrane protein design or modeling, we propose quantifying this vdW packing character of deeply engaging backbone “Holes” (<3.2 Å mainchain backbone contacts) as a novel structural quality metric to supplement or supplant current common ranking and filtering methods(8, 31). Support for this notion emerges from sidechain-to-backbone directed packing being a common feature of exceptionally stable TM protein complexes, including recent designs(7, 38, 59). In water, most forms of apolar burial are advantageous, even peripheral aliphatic contacts. By contrast, not just any apolar protein-protein contact arrangement will be more favorable than the chemically similar lipid-protein interactions competing within the membrane environment. Thus, vdW sterics of this form apparently optimized in Design-2 may be a general mechanism for how proteins achieve favorable apolar packing in membranes without the hydrophobic effect.

While A-X_6_-A motifs can achieve similar folds and stabilities as G-X_6_-G designs, glycines’ H-bonding adds another stabilizing force alongside vdW packing structures, exemplified by the strongest Design-2. Early work surmised that TM Cα H-bonds’ may contribute minimally and close Cα-H···O=C distances can simply be geometric consequences of more important interactions(60, 61). Design-2’s extensive close distanced H-bonding strongly points to the contrary – that Cα H-bonds bolster this TM structure. If the other G-X_6_-G designs can potentially achieve similar Cα H-bonding networks, then why are they less stable? We hypothesize the differences may arise from either weaker internal packing structures (less vdW surfaces from smaller or polar sidechains), not realizing similar strength Cα-H···O=C geometries, or an interrelated combination – i.e. not accommodating or balancing polar and packing features into their best forms, as in Design-2. Likewise, we are suspicious of whether interfaces of just two apolar helices lacking polar interactions, e.g. A-X_6_-A, can reach such heights of stability(62). However, the practical energy requirements for natural motifs are likely far less than these levels amongst optimized designs, especially when restrained by loops or domains. While TM spans in Nature typically suffice with short motif sub-segments (1-2 heptads like EmrE) often in kinked helices, design epitomizing those sub-structures into extended architectures yields highly stable and specific folding alongside idealized molecular features. Thus, our software effectively builds molecular archetypes of TM building blocks well suited for further deciphering of this and other membrane-specific motifs.

Advancing transmembrane-focused chemical principles and expanding the molecular design toolkit, alone or in combination with water-soluble engineering, will allow encoding of more sophisticated functions traversing different solvation environments such as sensing, cross-transport, tunable signaling(8, 30, 63–65). The effectiveness of the very simple 2-term generative design approach analyzing structure-aligned cross-evolutionary MSAs establishes that this depth and format of sequence-structure data is a viable starting point to train more generalized artificial intelligence models for improved prediction and design focused on membrane-spanning architectures.

## MATERIALS AND METHODS

### Protein design

Idealized homo-dimeric helices were modeled from parametric coiled-coiled equations (39) systematically varying inter-helix radius, α-helix frequency, super-coiling frequency (and interface or pitch angle), and z-offset. Using MASTER(40), Nine residue fragments of each helix from each model were queried for close geometric matches to non-redundant database membrane protein structures as of June 2020, prepared and curated at TM regions only as previously (38) (**Table S1**). Four unique starting backbones highly common were selected for *de novo* sequence design. The data-mining, statistical inference, and generative protein design algorithm are described in detail in the Extended Methods of the SI Appendix. MD simulations were prepared in Charmm-GUI(66) and run using GROMACS(67).

Structure prediction was done by ESMfold server(35) using TM sequence repeated twice bridged by 20x glycine linker or AF3 server (37).

### Expression and purification of designed TM proteins

SUMO-fusion constructs were cloned and expressed in pET-28a(+) vectors in C43(DE3) E. coli (Lucigen) grown in Terrific Broth (TB) with 50 µg/mL kanamycin induced with 0.5 mM IPTG at either 18, 30 or 37°C overnight, depending on the ideal temperature for that specific protein determined on a small scale. Cell were resuspended, lysed by tip sonicate, and extracted with 20 mM C14-Betaine (C14B, Sigma) detergent, prior to high speed centrifugate. Clarified lysate was purified by Nickle affinity (Ni-Indigo, Cube), eluted into buffer containing 8 mM C14B.

### Size-exclusion chromatography

Purified proteins were concentrated to 2-3 mg/mL and injected to a Superdex200i 10/300 column (Cytiva) at room temperature using a mobile phase of 25 mM NaPi at pH 8, 150 mM NaCl, 1 mM DTT, 0.5 mM EDTA and detergent (DDM, Anatrace; C14B; LDAO, Cayman).

### Gel electrophoresis

SDS-PAGE of Nickle-purified or SEC purified proteins were conducted either by the Biorad Tris/Glycine/SDS anyKd method or Invitrogen MES-SDS NuPage 4-12% method, which give different oligomerization banding or smearing behaviors. NuPage LDS sample buffer was mixed to final 1x concentration and ∼5 µg SUMO-TM protein is loaded, without boiling unless specified. Low SDS loading buffer was prepared as 0.2% SDS supplemented to native loading buffer: 10% glycerol, 50 mM Tris pH 6.8, 0.05% bromophenol blue final. TM peptides were first reconstituted to 1 mg/mL in 50 mM OG (Chem-Impex) or SDS before mixing with loading buffer.

### SARK-PAGE

Samples were mixed 1:1 with 0.2% or 2% w/v N-Lauroylsarcosine (Sark, Sigma) supplemented to native loading buffer, run on BioRad 12% polyacrylamide gels in Sark-Tris-Glycine running buffer (0.1% sarkosyl w/v, 25mM Tris pH 8.3, 192.5mM Glycine) as described previously (47).

### Glutaraldehyde cross-linking

Purified protein diluted to 0.1 mg/mL and pre-incubated with varied deregent concentrations was cross-linked with 5 mM glutaraldehyde (Sigma) for 1-60 minutes and quenched with 20 mM hydrazine monohydrate final (Sigma).

### Preparation of TM peptides

Cystine-labeled peptides were expressed as recombinant Sumo-fusion constructs as described above followed addition of thrombin protease (Sigma) added to sample prior to overnight dialysis in Tris buffer with 8 mM beta-mercaptoethanol. Minimal TM peptides for crystallography were digested overnight from SUMO fusion proteins by addition of porcine trypsin (1:100 mass ratio, Sigma). Cleaved TM peptides were purified HPLC using a Vydac C4 column with linear gradient of solvent A (99.9% water, 0.1% TFA) and solvent B (60% isopropanol, 29% acetonitrile, 9.9% trifluoroethanol, 1% water, 0.1% TFA). Identity and 95% purity were confirmed by LCMS (Waters)

### Thiol-disulfide equilibrium exchange

As previously described(42) for TM peptide glutathione redox equilibrium exchange 50µM each of N-terminal cysteine or C-terminal cysteine peptide was reconstituted in 100 molar excess detergent with 1.5mM [GSH]:[GSSG] (3:1 ratio) degassed redox buffer (25mM Tris, 200mM KCl, 0.5mM ethylenediaminetetraacetic acid (EDTA), pH 8.2) then incubated overnight to equilibration. Reaction products were analyzed by LC-MS with a UPLC C4 Column using a linear gradient of Solvent A and Solvent B (60% isopropanol, 35% acetonitrile, 4.95% water, 0.05% TFA).

### **X-** ray crystallography

Design-2 TM peptide was crystallized by hanging drop vapor diffusion in 0.1 M Sodium chloride, 0.1 M Lithium sulfate, 0.1 M Sodium citrate pH 3.5, 30% PEG400, flash frozen without additional cryoprotectant, and diffracted at APS 23-ID-B. The design model for molecular replacement in Phenix(68).

### Quantum Mechanics (QM) calculations

The coordinates for the two glycines (G16 from chain A and chain B of pdb 8SRN) and backbone atoms from adjacent residues to yield an N-terminal acetyl cap and C-terminal methyl capped tertiary fragment subjected to constrained minimization at B3LYP/6-31G* level of theory with GD3 empirical dispersion using Gaussian09(69) with a harmonic potential of 0.0005 Hartree/Bohr(70) applied towards the initial conformation. The resulting wave functions were analyzed using Multiwfn(70).

### Structural informatics analysis of membrane protein structural database

Cα-H···O=C hydrogen bond interactions were identified using the geometric criteria described by Senes et al (4) searching unique instances (accounting for symmetry) in non-redundant membrane spanning domains (**Table S1**), listed in Dataset S1. Each Cα proton was assigned as R or S.

## Data availability

The X-ray structure of G-X6-G Design-2 has been deposited to the Protein Data Bank with code 8SRN. The source data for biophysical characterization, molecular models, and uncropped gels have been uploaded to a zenodo data repository with DOI: 10.5281/zenodo.15476293

## Code availability

The code for sequence design is hosted at https://github.com/JayGolden55/tmDimer

## Acknowledgements

The authors thank the support of Robyn Stanfield and Ian Wilson in conducting our crystallography work. We thank Huong Kratochvil for careful reading of the manuscript and insightful comments. M.H. & S.F. are supported by R01GM069832. C.A.M. was supported by the Diekman Family Graduate Fellowship and ARCS Fellowship. C.A. was supported by the UCSD McNair Scholars Program. Use of the Stanford Synchrotron Radiation Lightsource, SLAC National Accelerator Laboratory, is supported by the U.S. Department of Energy, Office of Science, Office of Basic Energy Sciences under Contract No. DE-AC02-76SF00515. The SSRL Structural Molecular Biology Program is supported by the DOE Office of Biological and Environmental Research, and by the National Institutes of Health, National Institute of General Medical Sciences (P30GM133894).

## Appendix of of Supplementary Materials

**- Supplementary** Figures 1-16

**- Supplementary Tables 1 -5**

**- Extended Data Table 1**

---

**Supplementary Figure 1.**
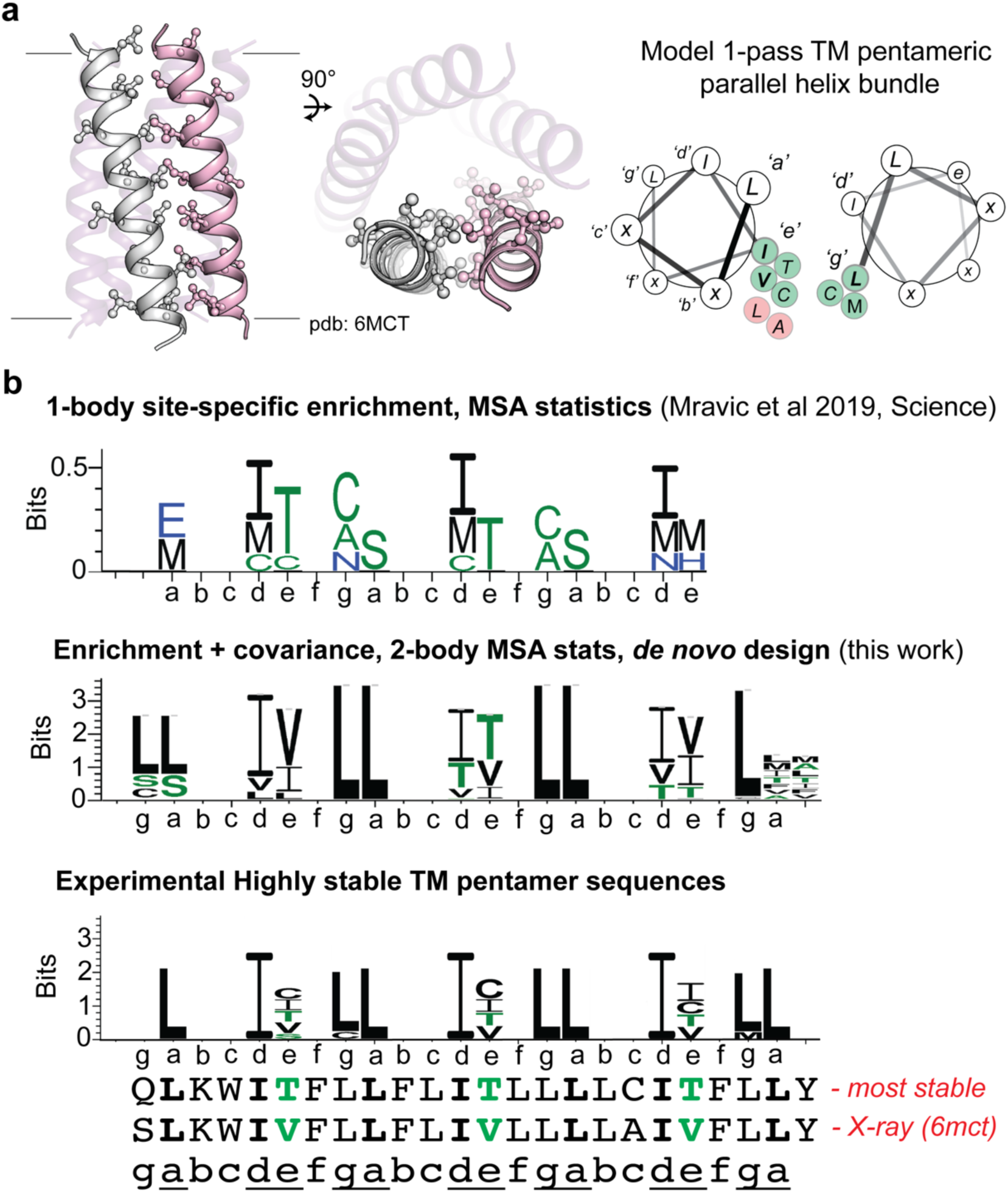
Benchmark of multi-body sequence statistics generative design algorithm on family of synthetic pentameric proteins with a common parallel TM building block. **(a)** Structural model of single-span (1-pass) pentameric protein “eVgL” (pdb: 6mct) with sidechain sticks representing its apolar sidechain steric packing motif, left. Right, helical wheel diagram with heptad assignment and sequences used in the steric packing pattern for this TM building block. Amino acids in green at ‘g’ and ‘e’ positions are those experimentally tested to lead to highly stable folding. **(b)** Top, Weblogo of site-specific amino acid enrichment per residue when analyzing log odds ratios of amino acids frequencies occurring in similar helix-helix geometries in natural proteins, relative to the background distribution (expected frequencies) of amino acids in membrane proteins, adapted from ref ^2^ (Mravic et al, 2019, Science). Middle, sequences from the generative sequence design, combining inter-residue covariance (2-body) with site-specific enrichment (1-body) statistics. Bottom, interface residues in experimentally tested variants of the model parallel single-span pentameric protein varying at ‘e’ and ‘g’ positions. Full sequences of eVgL and eTgL, the latter being the most stable variant, with ‘e’ position in green and aligned by heptad position; interface residues underlined.

**Supplementary Figure 2.**
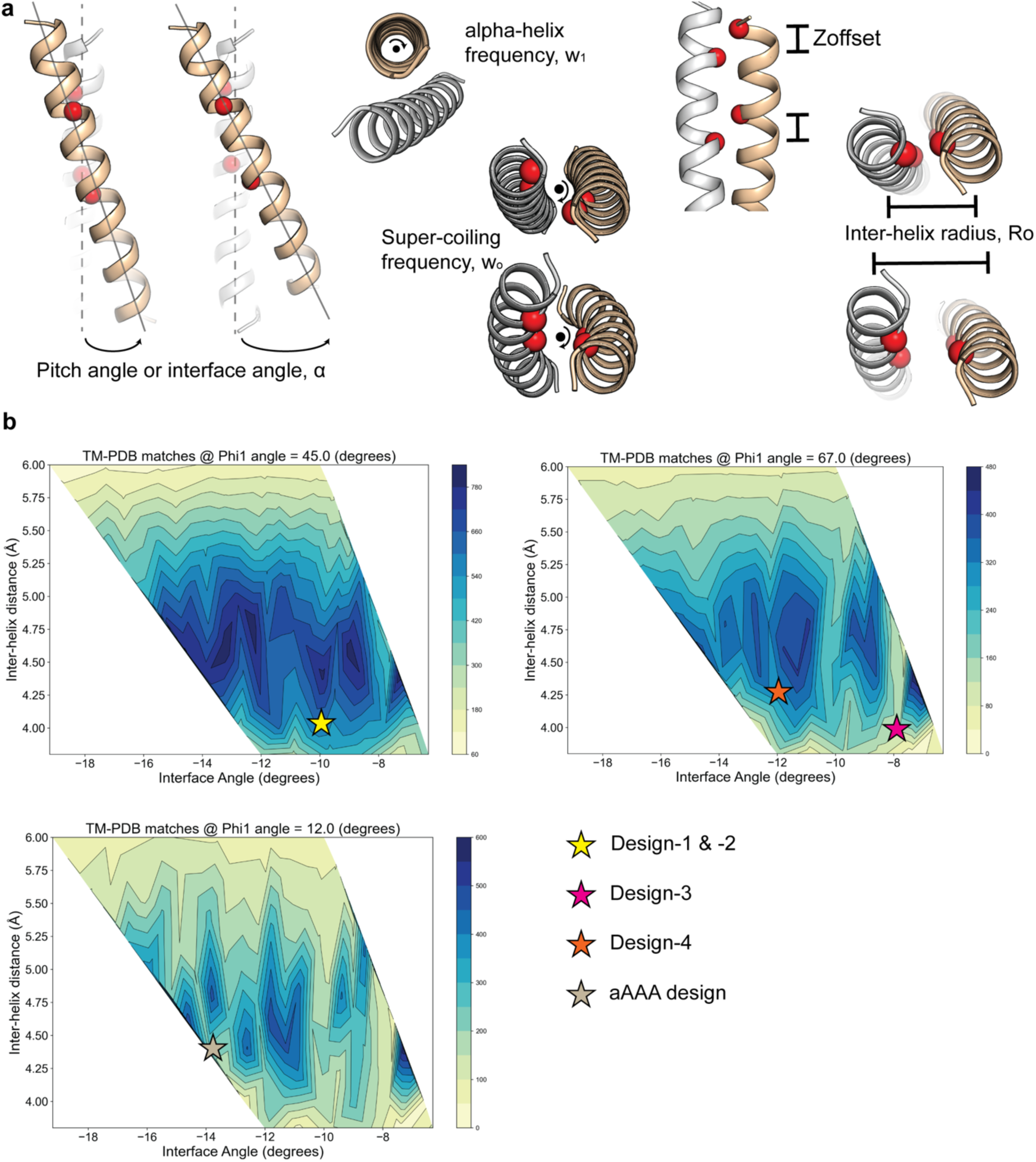
Coiled-coil backbone geometries sampled and designs for antiparallel TM helices. **(a)** Helix-helix interactions in 2 transmembrane spans approximated by parametric coiled coils, based on the common building blocks previously observed for antiparallel interacting helices in membrane proteins. Pitch angle or interface angle, α; alpha-helix frequency ω_1_ internal rotation; super-helical or super-coiling frequency ω_0_ around coiled-coil axis; Z-offset between helices or helix register; inter-helical radius. **(b)** Heatmap of number of close geometric matches (1 Å backbone RMSD) for representative 12 residue fragments of each *de novo* parametric coiled-coil model searched into a database of non-redundant membrane protein structures (Table S5), taken as the average of models whose Z-offsets range. Plots (top left, top right, bottom left), shown for discrete alpha-helix register aka internal rotation (ω_1_). The geometry of each designed TM protein starting models is plotted as colored stars.

**Supplementary Figure 3.**
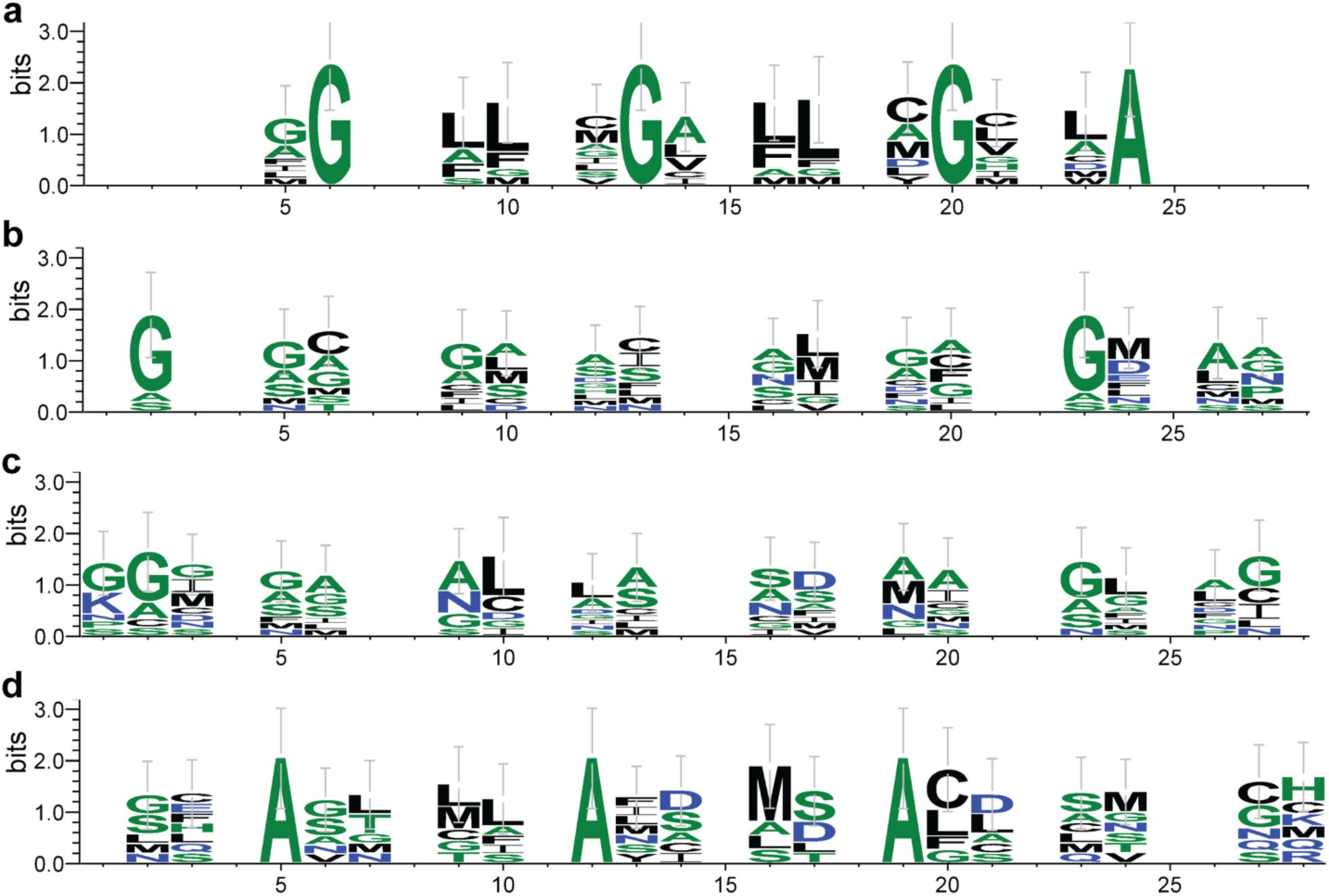
Sequence logos of top designs for each antiparallel TM helix backbone dimer model. Weblogo of sequences generated by protein design for each backbone structure in tested designs. (**a**) Logo of G-X_6_-G Design-1 and −2 backbone, top 10 sequences (**b**) Sequence logo of G-X_6_-G Design-3, top 10 sequences (**c**) Sequence logo of G-X_6_-G Design-4, top 10 sequences (**d**) Sequence logo of A-X_6_-A Design backbone, top 7 sequences

**Supplementary Figure 4.**
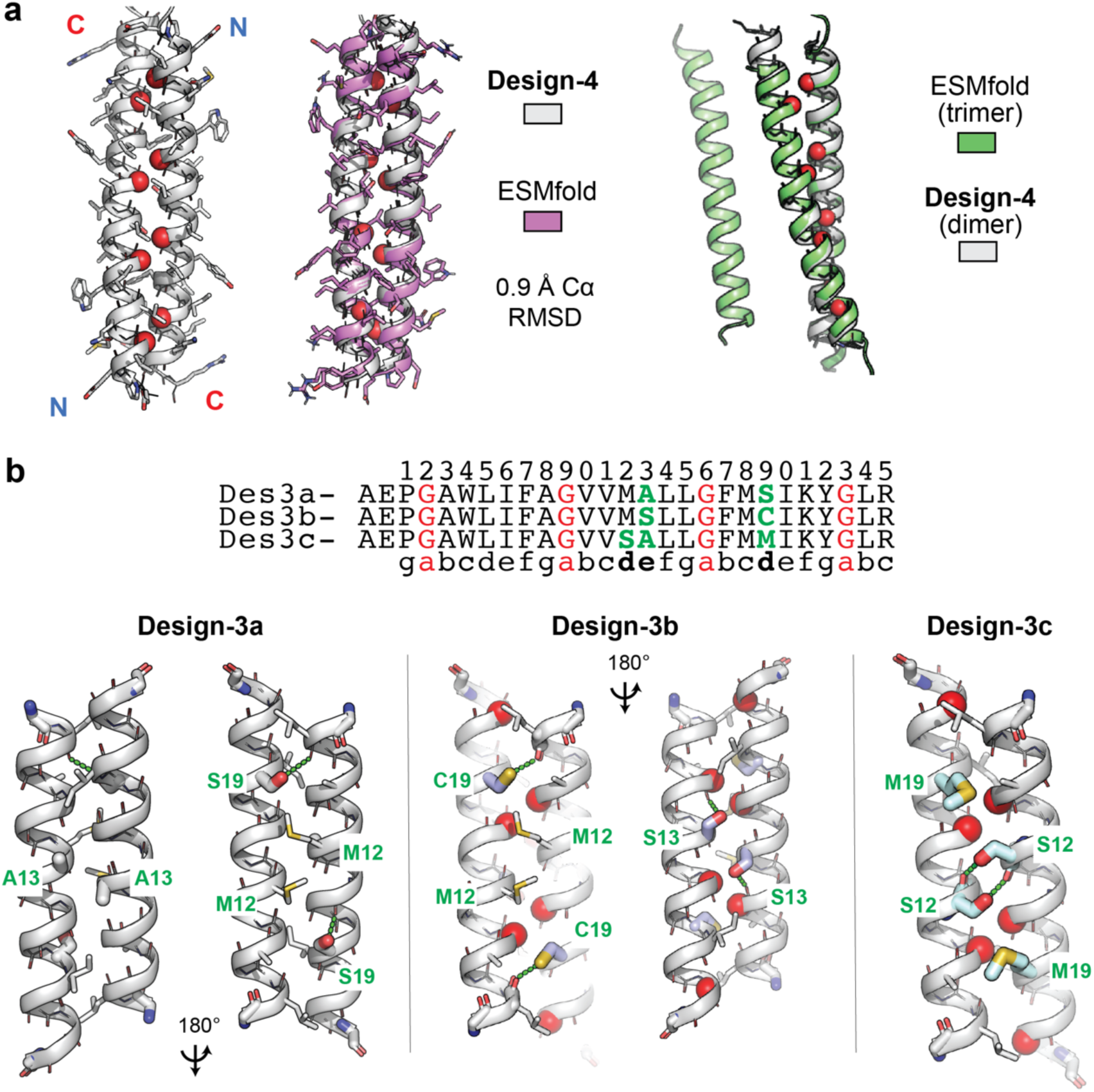
Ab inito structure prediction of G-X6-G de novo TM domain Design-4 and Design-3. (**a**) Top, alignment of Design-3 variants labeled by 7-residue heptad repeat differing at key ‘d’ and ‘e’ positions (bold heptad label, green amino acids). Bottom, structural models predicted for Design-3 variants a-c, with key differing amino acids labelled in green; glycines, red spheres. Hydrogen bonding across the helix are expected at Cys/Ser19 and Ser12, but not Ser13. (**b**) Left, ESMfold predicted structure (magenta) versus design model (white) for dimeric G-X_6_-G Design-4, matching at <1 Å RMSD. Motif glycines in red. Right, predicted trimer model (green) for the Design-4 sequence results in 2 helices adopting the same antiparallel dimer conformation as the design (white) with the additional TM span predicted to not interact.

**Supplementary Figure 5.**
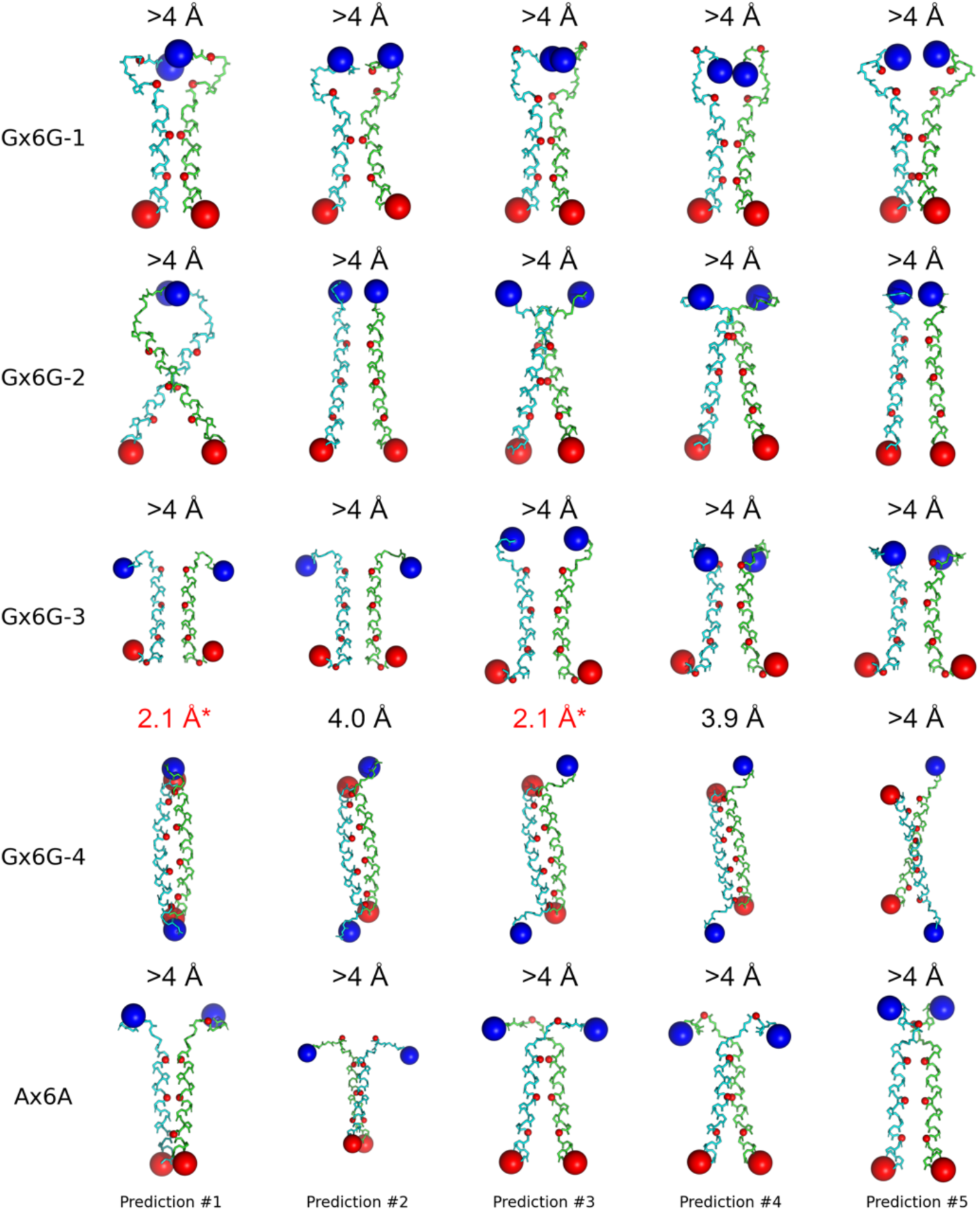
AlphaFold-3 predictions of G-X_6_-G and A-X_6_-A designed TM domain homo-dimers. Top 5 predictions of each TM protein design homo-dimer (label as “GX6G-1” = Design-1) shown as cyan or green mainchain ribbons colored by unique chain, and N-termini and C-termini colored as blue and red spheres respectively. C-alpha atom RMSD is listed for the best alignment of each prediction to the 200 ns MD simulation frame design model or X-ray structure (G-X_6_-G Design-2). The design with the longest hydrophobic TM span G-X_6_-G Design-4 is predicted as antiparallel matching the design model for predictions 3, 4, and 5. For the other designs, all AF3 predicted structures are parallel and incorrect relative to the antiparallel structure observed by crystallography and ESMfold predictions.

**Supplementary Figure 6.**
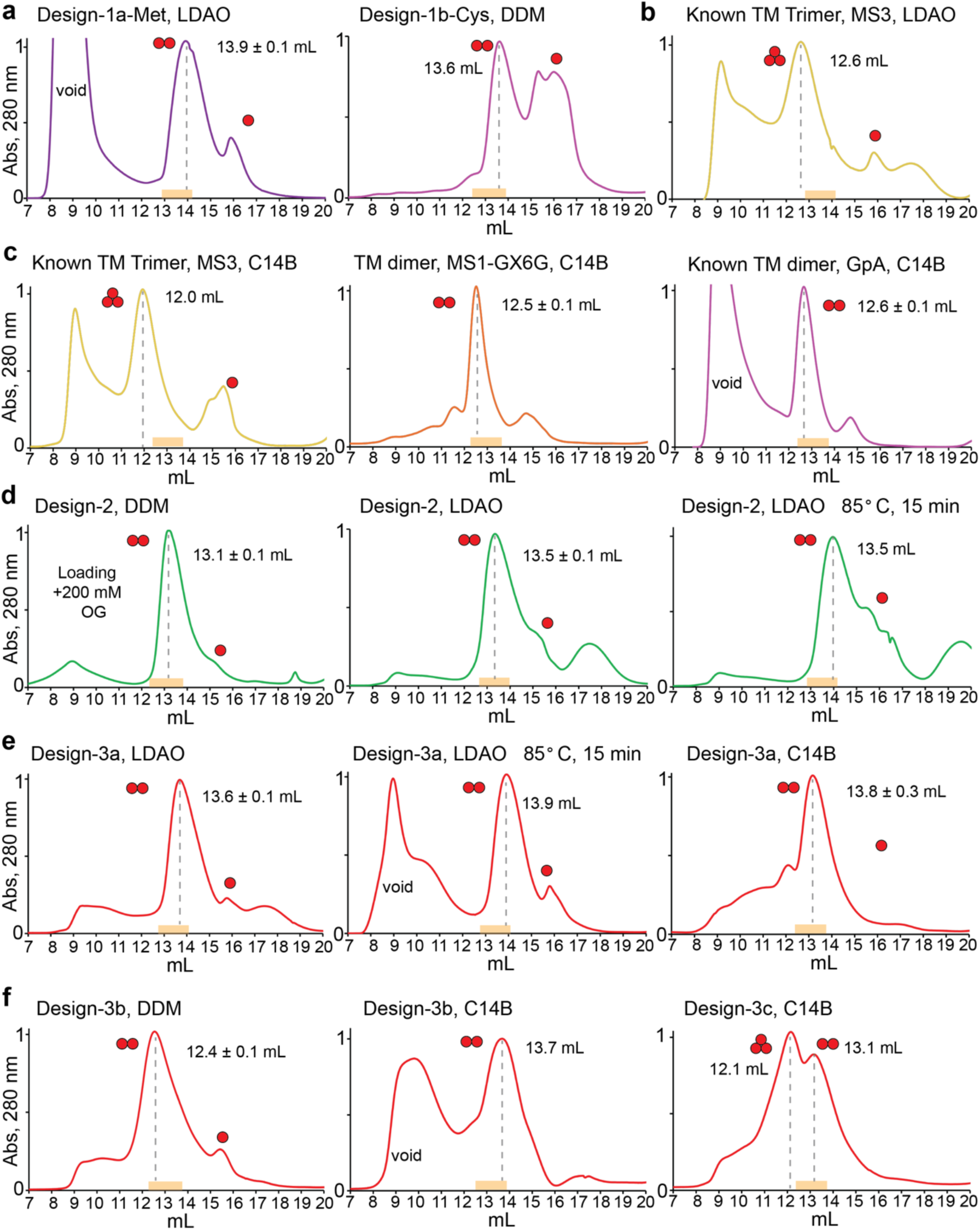
Size Exclusion Chromatography Folding Test of SUMO-TM proteins. **(a)** Design-1a and 1b in LDAO (n=3) or DDM (n=2) mobile phases, respectively**. (b)** Reference TM domain trimer, MS3 in LDAO (n=1). **(c)** Reference TM domain trimer, MS3, and two dimers, MS1-GX6G and GpA, in C14B, representative of n=3 experiments. **(d)** Design-2 in DDM and LDAO (each n=3), including in LDAO with 15-minute pre-incubation at 85° C (n=1). **(e)** Design-3a in C14B and LDAO (each n=3), including in LDAO with 15-minute pre-incubation at 85° C (n=1). **(f)** Left, Design-3b in DDM and C14B (each n=3). Right, Design-3c in C14B elutes as two peaks (n=1). Chromatograms are 280 nm absorbance (normalized to peak max) from 2-3 mg/ml protein in superdex200i 10/300 column, mobile phase: 50 mM Na Phosphate pH 7.8, 100 mM NaCl, 0.5 mM EDTA, 1 mM DTT, and detergent (6 mM LDAO, 1 mM DDM, or 3 mM C14B). Red spheres, assigned oligomer.

**Supplementary Figure 7.**
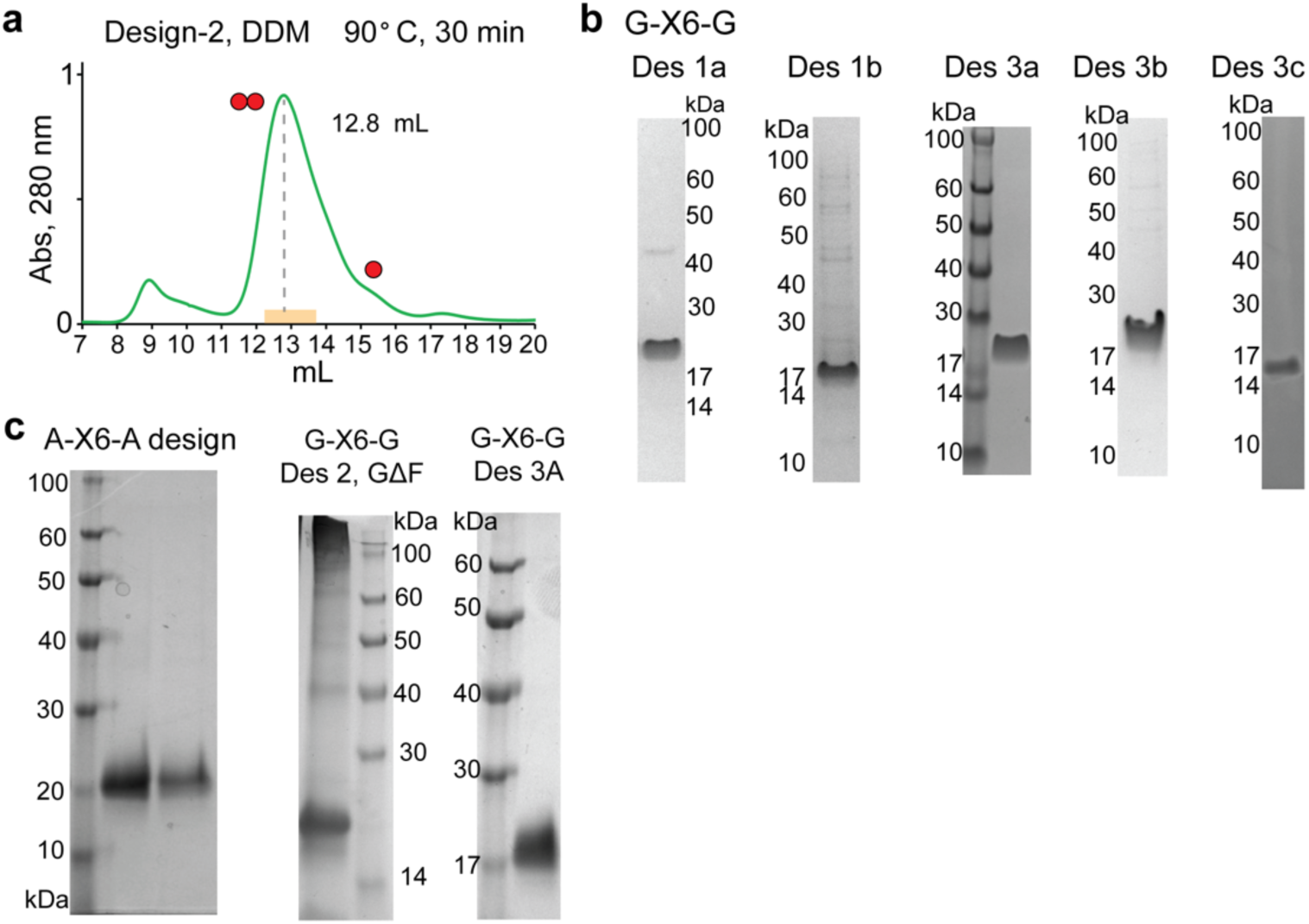
Additional characterization of SUMO fusion TM designed antiparallel oligomers. (**a**) Design-2 at 2 mg/mL after 30-minute incubation at 90° C run in a superdex200i 10/300 column at 25° C, mobile phase: 50 mM Na Phosphate pH 7.8, 100 mM NaCl, 0.5 mM EDTA, 1 mM DTT, and 1 mM DDM. Chromatograms is 280 nm absorbance (normalized to peak max), representative of n=3 trials. Single and double red circles denote monomer (∼15.5 mL) and dimer (12.8 mL) peaks. Orange rectangle denotes expected dimer range for a single-span SUMO-TM fusion in DDM. (**b**) Purified SUMO fusion TM domain constructs for Designs 1a, 1b, 3a, 3b, and 3c in NuPage Bis-Tris gel ran with MES-SDS running buffer, representative of n=3 trials. (**c**) Purified SUMO fusion TM domain constructs for the A-X_6_-A design (left), mutant G-X_6_-G Design-2 GΔF (middle), and G-X_6_-G Design-3a in BioRad anyKd gel ran with Tris-Glycine-SDS running buffer. All proteins run as predominantly monomers under these conditions, representative of n=3 trials.

**Supplementary Figure 8.**
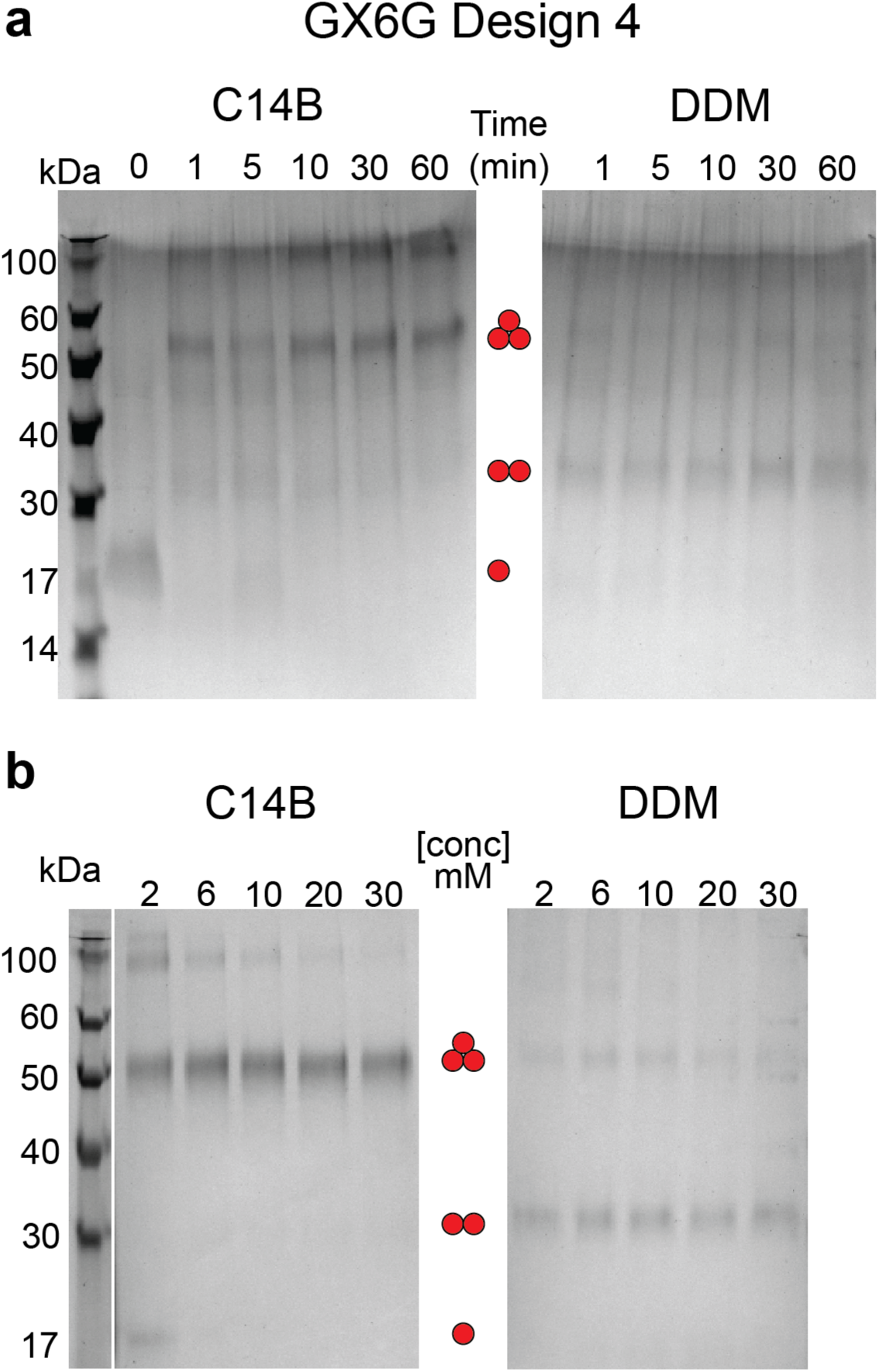
Glutaraldehyde cross-linking of Design-4. **(a)** 2 mM glutaraldehyde was used to cross-link SUMO-Design-4 (17 kDa) in C14B (left) or DDM (right) for 1 to 60 minutes, resulting in rapid formation of primary trimers in C14B and dimers in DDM (n=1 experiment). Red circles denote monomer, dimer, trimer peaks. Buffer is the mobile phase from SEC with 0.1 mg/mL total protein in the reaction. **(b)** Gluteraldehyde cross-linking for 10 minutes after pre-incubating the protein in a titration of detergent concentration to alter the proteins dilution in the micelle phase (n=1 experiment). All conditions form the primer covalent oligomer as in panel a, with reduced amount of higher-order species at higher detergent concentrations.

**Supplementary Figure 9.**
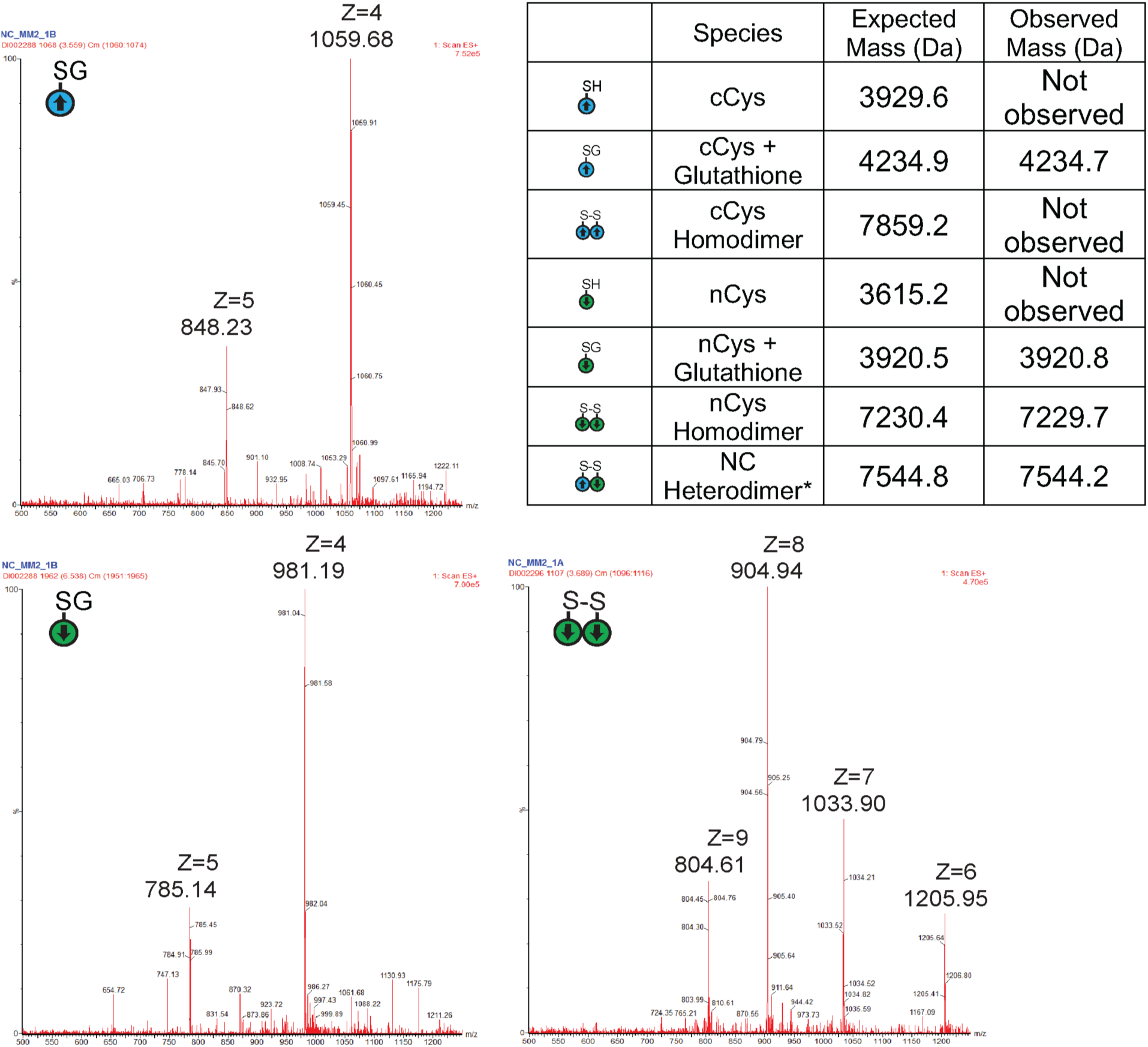
Mass Spectrometry of Design-1a TM peptides in thiol-disulfide exchange. Table of all expected and observed N-cys and C-Cys TM peptide species upon reversible glutathione oxidation. Representative (n=3) mass spec identification of the main Design-1a monomer, glutathione adduct, and covalent dimer species in the LC trace in Figure 3. Mass spectra are representative of n=3 experiments

**Supplementary Figure 10.**
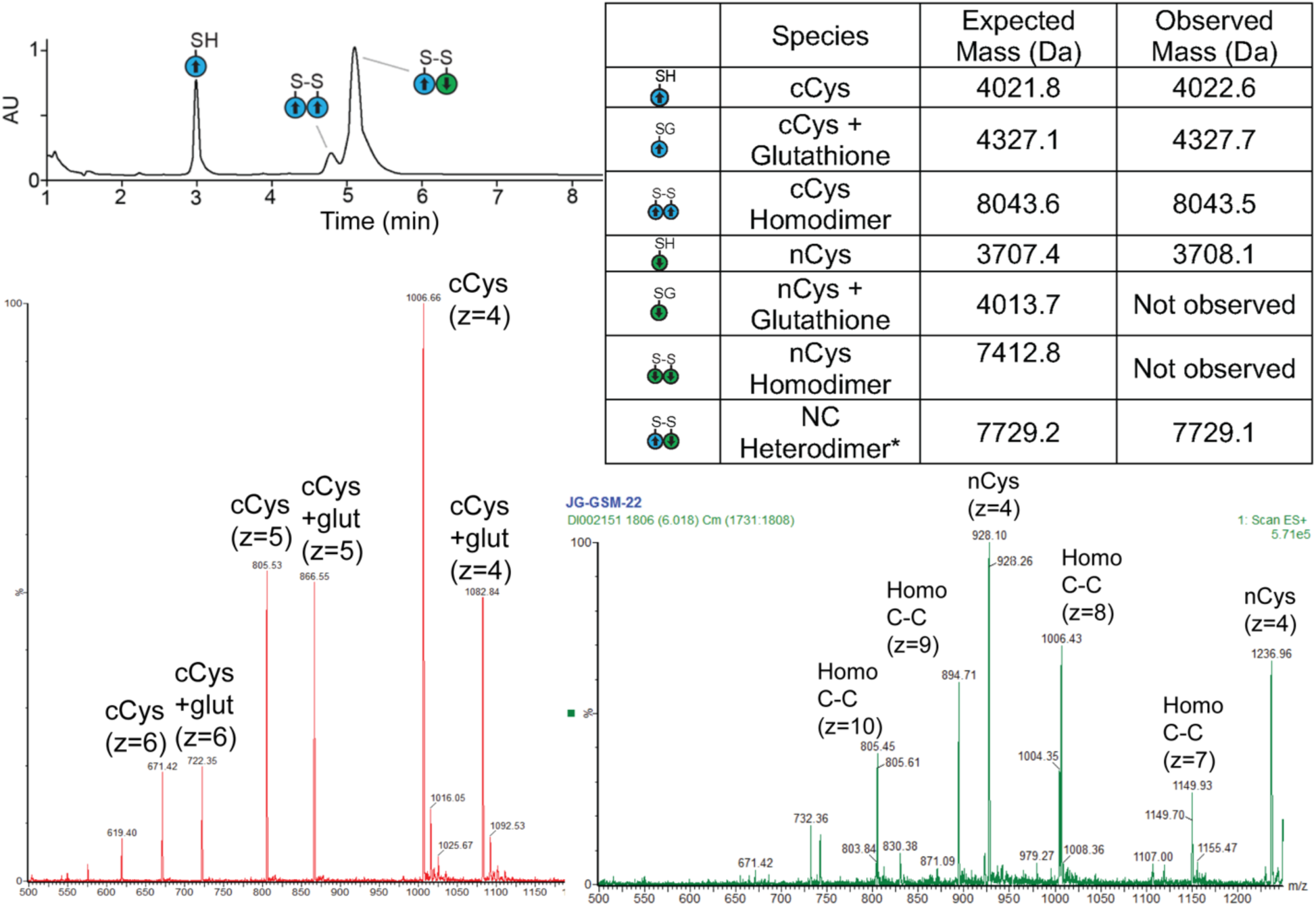
Mass Spectrometry of Design-2 TM peptides in thiol-disulfide exchange. Top left, LC trace with cartoons of covalent monomer or dimer species as in Figure 3a. Top, right, table of all expected and observed nCys and cCys TM peptide species upon reversible glutathione oxidation. Representative mass spec identification of the main Design-2 monomer, glutathione adduct, and covalent dimer species in the LC trace in Figure 3. Bottom Left, mass spectra at ∼ 3 min peak, containing mixture of monomeric cCys and its glutathione adduct. Bottom right, minor front peak at ∼4.8 min containing mix of nCys monomer and parallel homodimer of cCys species (Homo C-C). UV trace is max plot, condensation of all UV wavelengths in absorbance units (AU). All mass spectra are representative of n=3 experiments

**Supplementary Figure 11.**
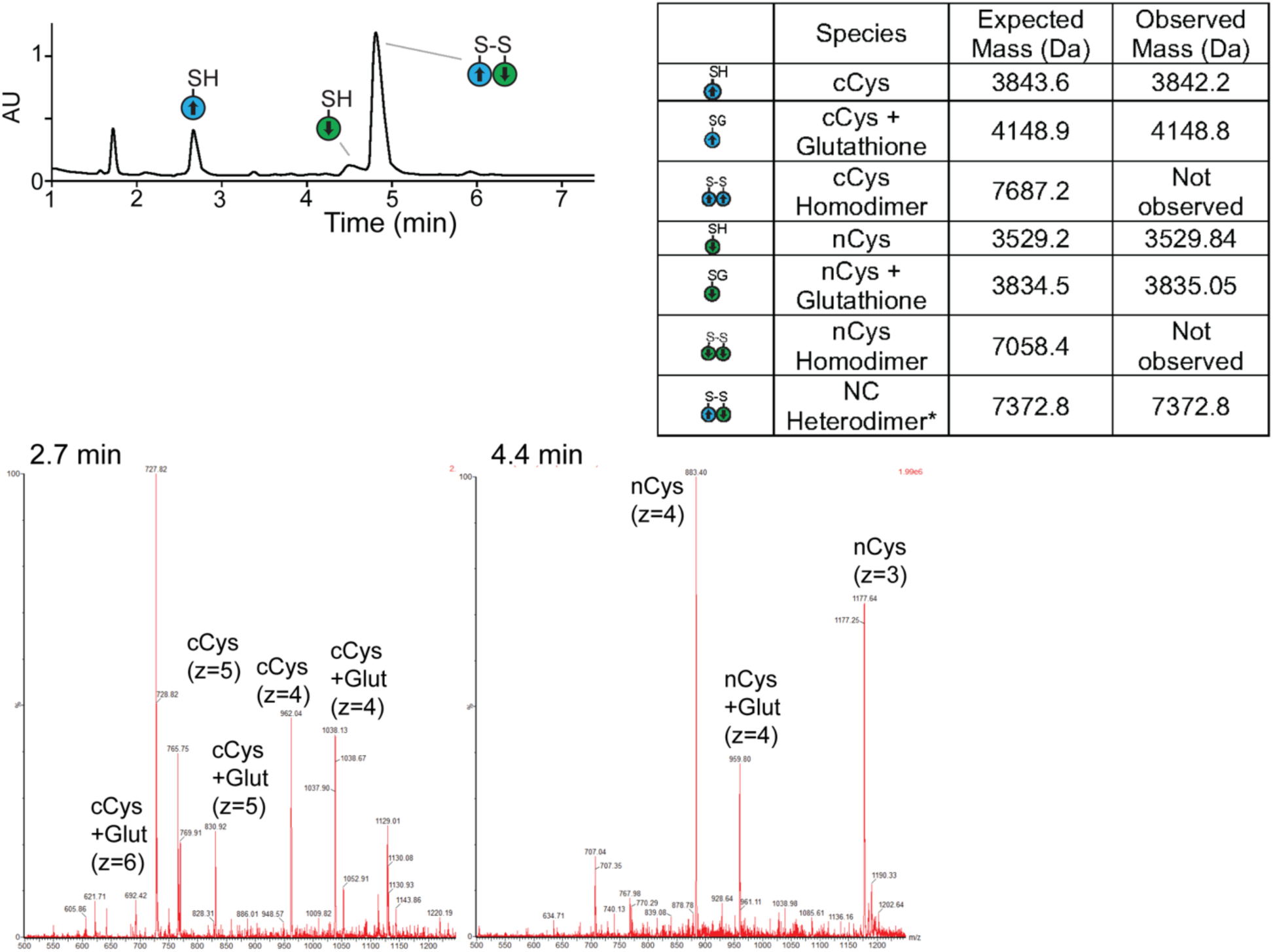
Mass Spectrometry of Design-3a TM peptides in thiol-disulfide exchange. Top left, LC trace with cartoons of covalent monomer or dimer species as in Figure 3a. Top, right, table of all expected and observed nCys and cCys TM peptide species upon reversible glutathione oxidation. Representative mass spec identification of the Design-3a species eluted in Figure 3. Bottom Left, mass spectra at ∼2.7 min peak, containing mixture of monomeric cCys and its glutathione adduct. Bottom right, minor front peak at ∼4.4 min containing mix of nCys monomer and its glutathione adduct. The UV peak at 1.8 min did not contain peptide material in its mass specta. UV trace is max plot, condensation of all UV wavelengths in absorbance units (AU). Mass spectra are representative of n=3 experiments

**Supplementary Figure 12.**
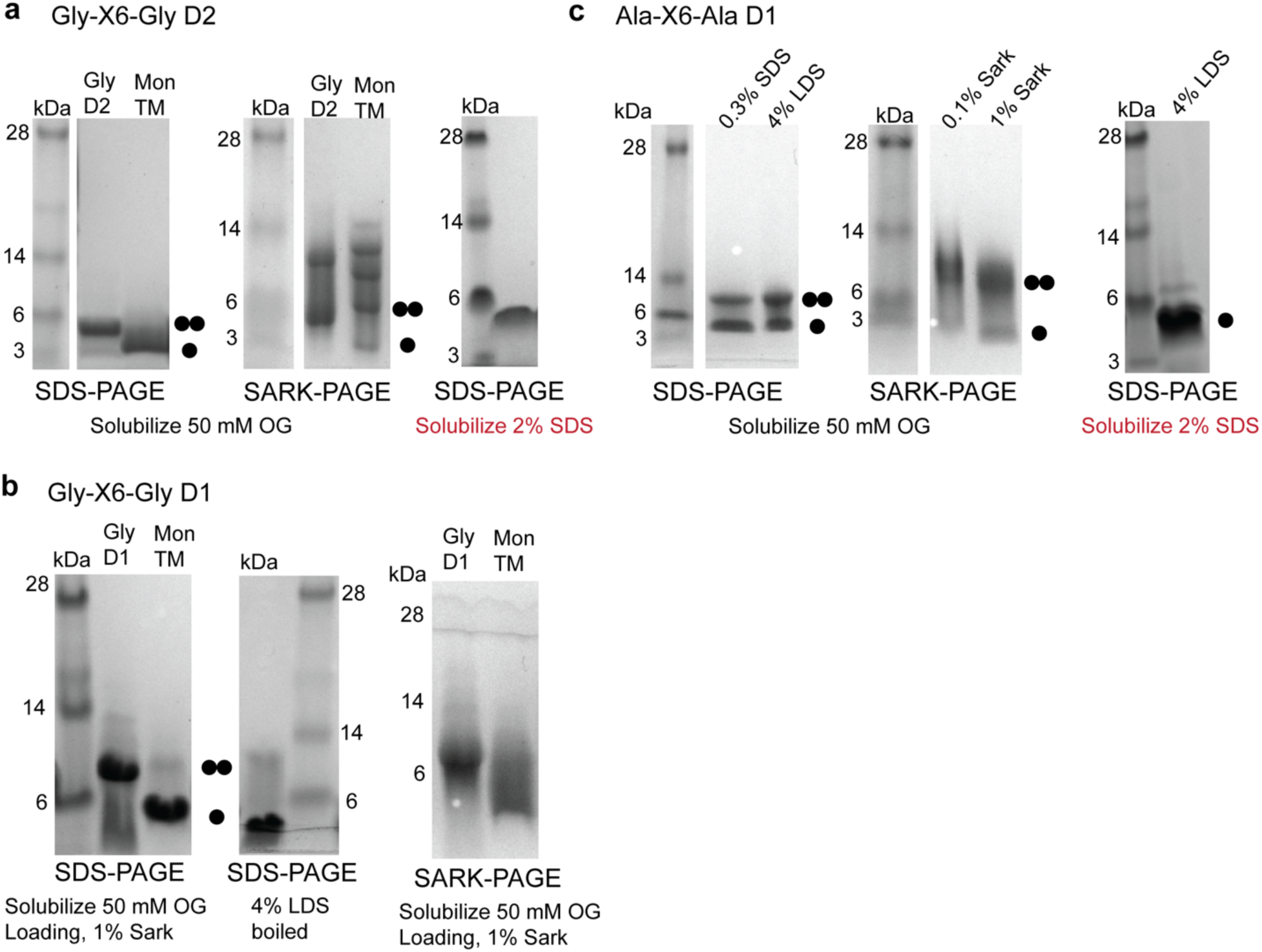
SDS and sarkosyl (SARK) PAGE of minimal TM peptide designs. **(a)** G-X_6_-G Design-2 (D2) minimal TM peptide gel migration. Left, Design-2 TM peptide solubilized to 2 mg/mL in 50 mM OG migrates as two possible bands with major band at just below 6 kDa and lighter possible band at 3 kDa, compared to a reference known monomer TM peptide migrating at 3 kDa. Middle, in the more gentle SARK-PAGE, the Design-2 runs asa mix of the 6 kDa species and a higher species ∼10 kDa. Non-specific laddering of the monomer for comparison. Right, Design-2 solubilized in 2% w/v SDS run in SDS-PAGE migrates as a single species ∼5 kDa. These data are unclear if the ∼5 kDa species is the SDS-resistant dimer or a monomer, but overall the protein exhibits oligomerization propensity, all representative of n=2 trials. **(b)** G-X_6_-G Design-1a (D1) TM peptide migrates as two species, above 6 kDa and below 6 kDa, which is destabilized to the lower molecular weight species, likely monomer, upon boiling and loading in high LDS detergent (left and middle, representative of n=2 trials, compared to monomeric TM peptide). Right, SARK-PAGE results in the >6 kDa species being stabilized, using 1 % SARK loading buffer. **(c)** A-X_6_-A designed TM peptide migrates as two clear species in SDS-PAGE solubilized in OG with minimal SDS loading buffer (0.3 w/v%) or harsher 4% LDS, above and below 6 kDa similarly to the G-X6-G designs (left, representative of n=3 trials). Middle, the same two bands appear in SARK-PAGE with the sample in 0.1% or 1% SARK loading buffer. Right, when the peptide are instead originally solubilized in 2% SDS, mixed with 4 % LDS loading buffer and ran on SDS-PAGE, it runs as the lower < 6 kDa molecular weight species.

**Supplementary Figure 13.**
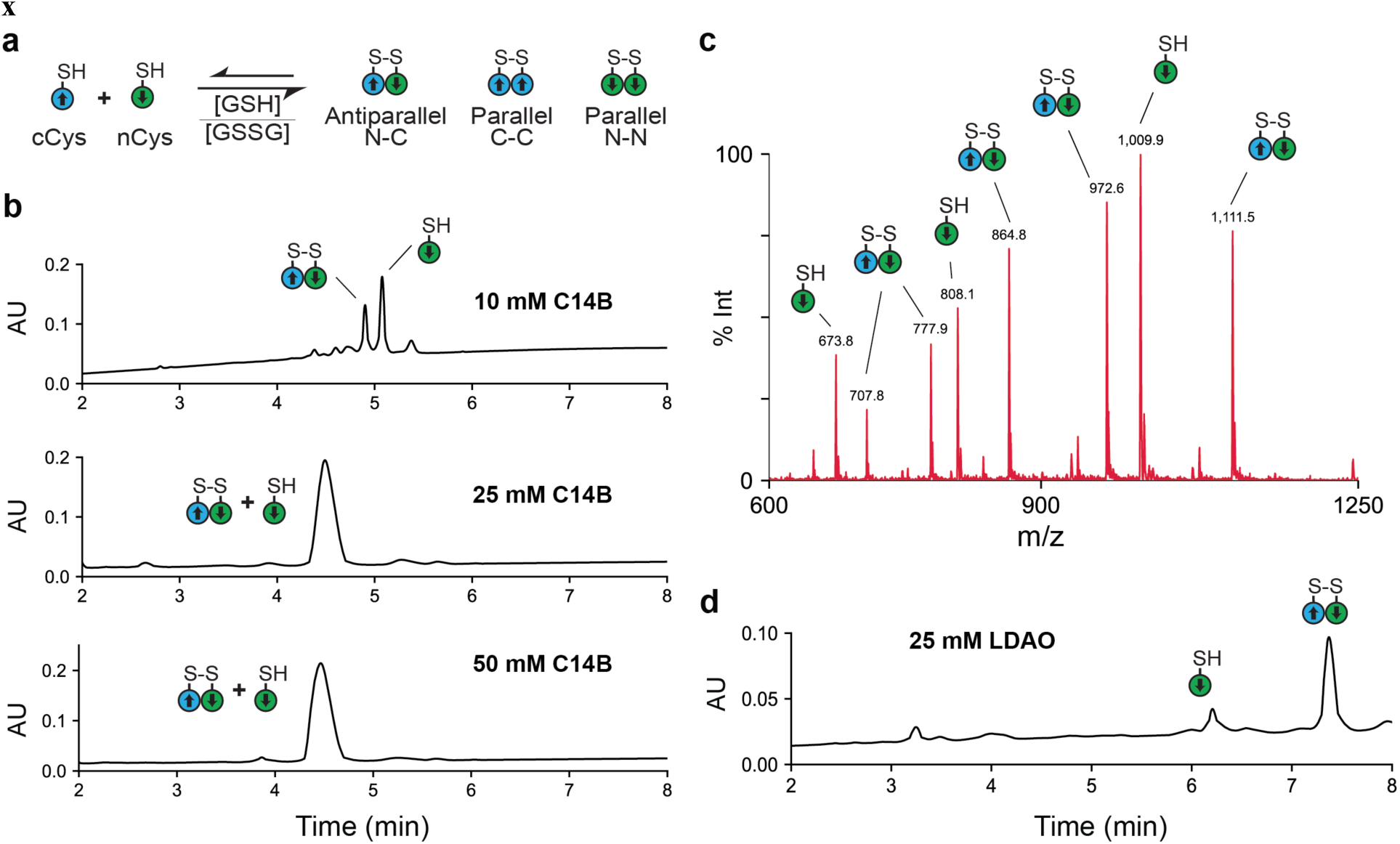
Thiol-disulfide exchange and mass spectrometry of A-X_6_-A TM peptides. (**a**) Scheme of thiol-disulfide equilibrium exchange of nCys and cCys TM peptides. (**b**) UV max plot trace in absorbance units (AU) of thiol-disulfide reaction of A-X_6_-A peptide mixture shows two main peaks of protein material when reconstituted in 10, 25, or 50 mM C14B detergent (representative of n=3). The mass spectra shows overlapping elution of the Ncys monomer and N-C antiparallel homodimer. Thus, the N-C heterodimer is the prominent disulfide bonded species observed above the detection limit. (**c**) Mass spectra of main peak in panel (b), identifying the N-C heterodimer (expected: 7771.418 Da; observed 7772.86 Da) and glutathione adduct of the Ncys monomer (expected: 4034.8 Da; observed 4036.0 Da). (**d**) UV max plot trace in absorbance units (AU) of thiol-disulfide reaction of A-X6-A peptide mixture when reconstituted in 25mM LDAO detergent shows that the major UV peak is the N-C antiparallel homodimer, and it is the only disulfide bonded species detectable (representative of n=3).

**Supplementary Figure 14.**
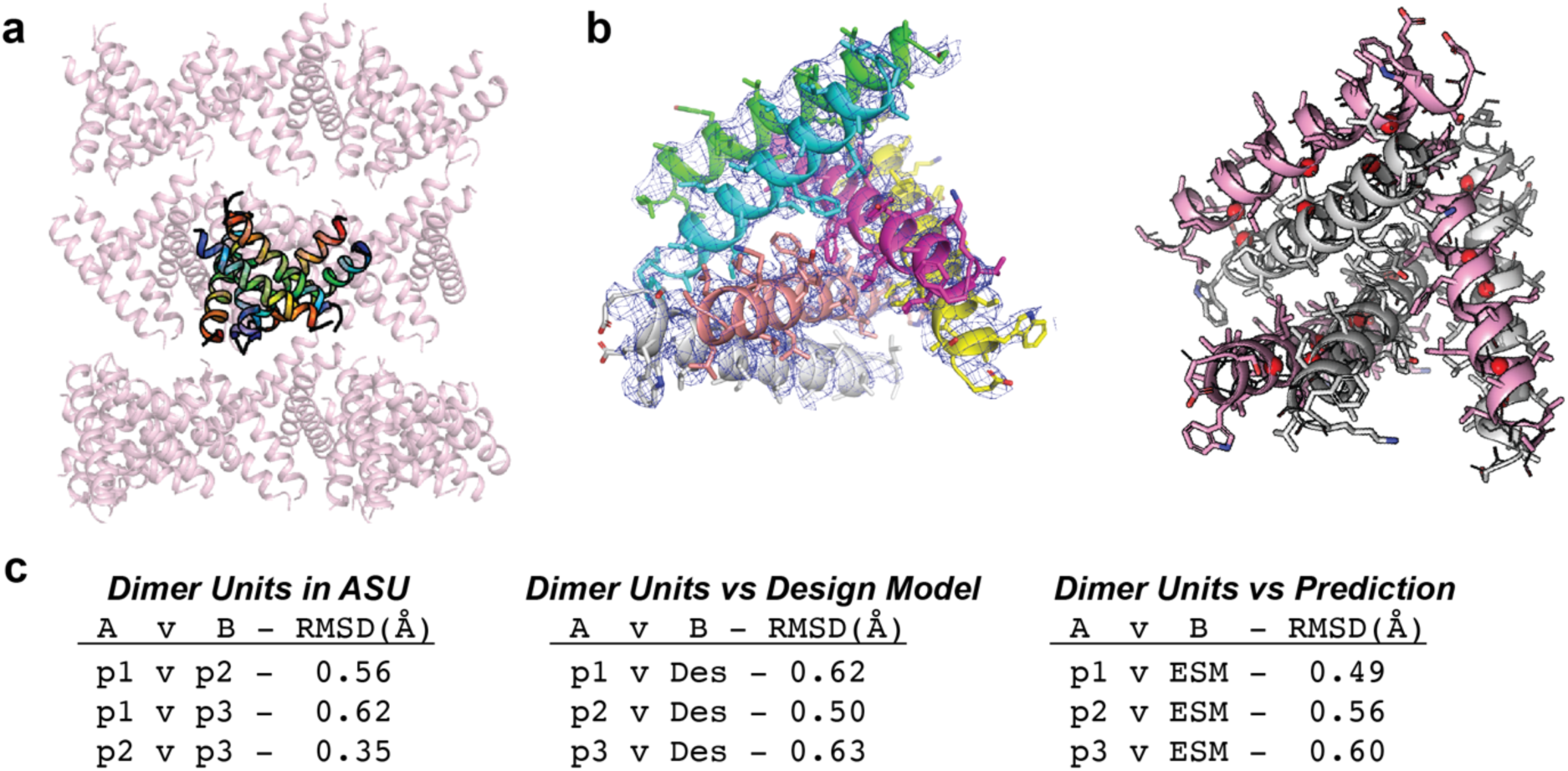
Design-2 TM peptide crystal structure. (**a**) Crystal packing in layers with asymmetric unit (ASU) of 6 protomers in chainbows and symmetry related molecules in pink. (**b**) Left, 2Fo-Fc electron density contoured at 1 sigma for the Design-2 ASU colored by chain. Right, the three pseudo-identical antiparallel dimer units colored as white and pink pairs, with glycine residues as red spheres. (**c**) Tables of backbone RMSD. Left, RMSD between the 3 antiparallel dimer units (p1, p2, p3) and each other. Middle, RMSD of dimers versus the intended design model. Right, RMSD of dimers versus the independent model from ESMfold structure prediction.

**Supplementary Figure 15.**
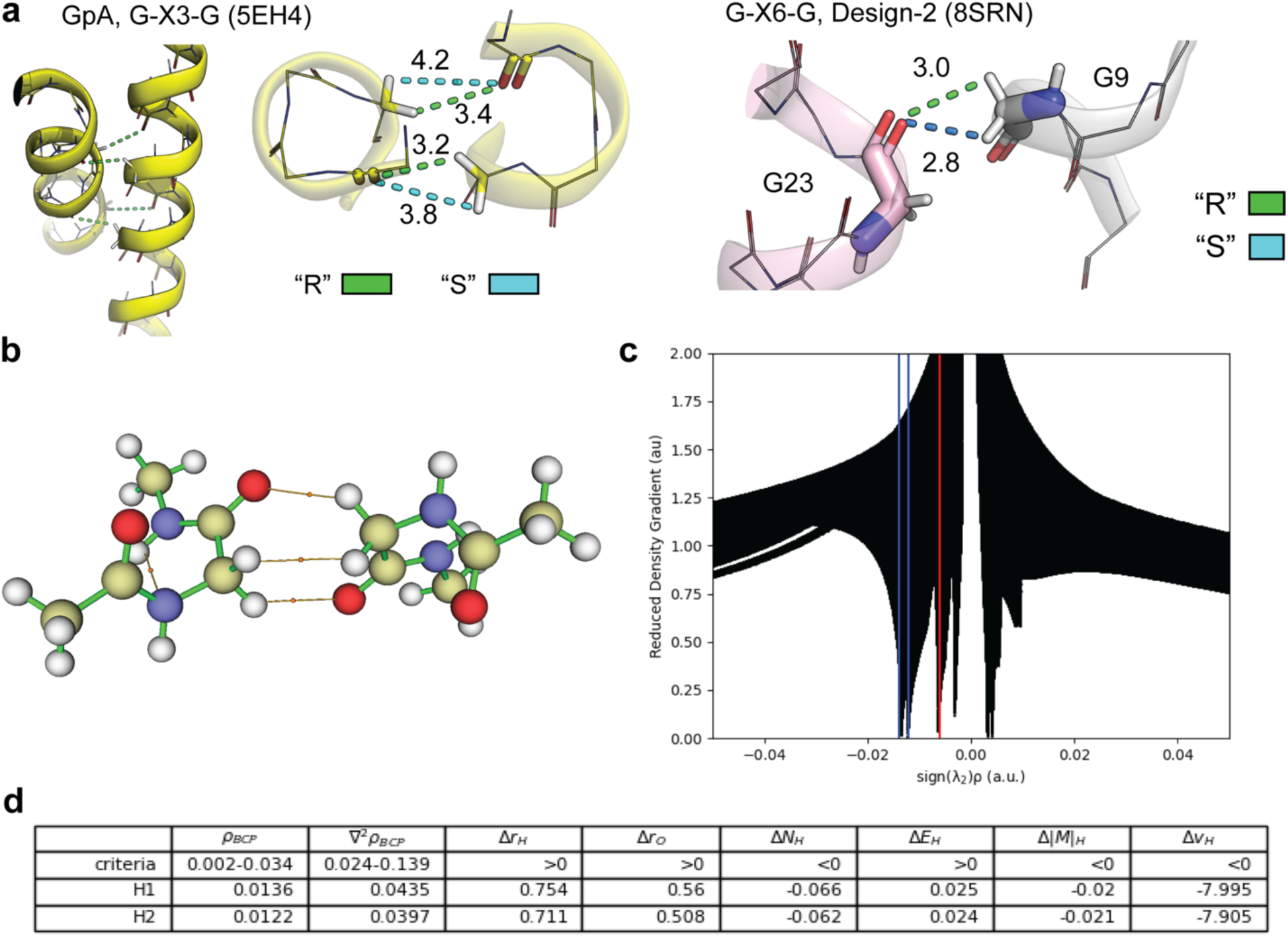
Cα H-bonding analysis G-X_6_-G motifs in natural and designed membrane proteins. (**a**) Left, pseudo-chrial R proton (green dash) forming Cα H-bonds in Glycophorin A LCP crystal structure, with S protons being too far for H-bonding (blue). Right, antiparallel Cα H-bonds in G-X_6_-G de novo design TM protein (pdb: 8SRN) which much closer distances. (**b**) Molecular graph of the QM optimized glycine interface. Bond critical points shown as small orange circles (**c**) Reduced density gradient NCI s vs. sign(λ2)ρ plot of the glycine interface, calculated at the di-glycine interface. C-H…O hydrogen bonds marked in blue, C-H…H-C vdW interaction marked in red (**d**) AIM properties of the studied Cα-H…O bonds (in a.u.) from the QM minimized G-X6-G glycine packing motif. Criteria for hydrogen bond adapted from Koch & Popelier (Citation 56 in main text).

**Supplementary Table 1.**
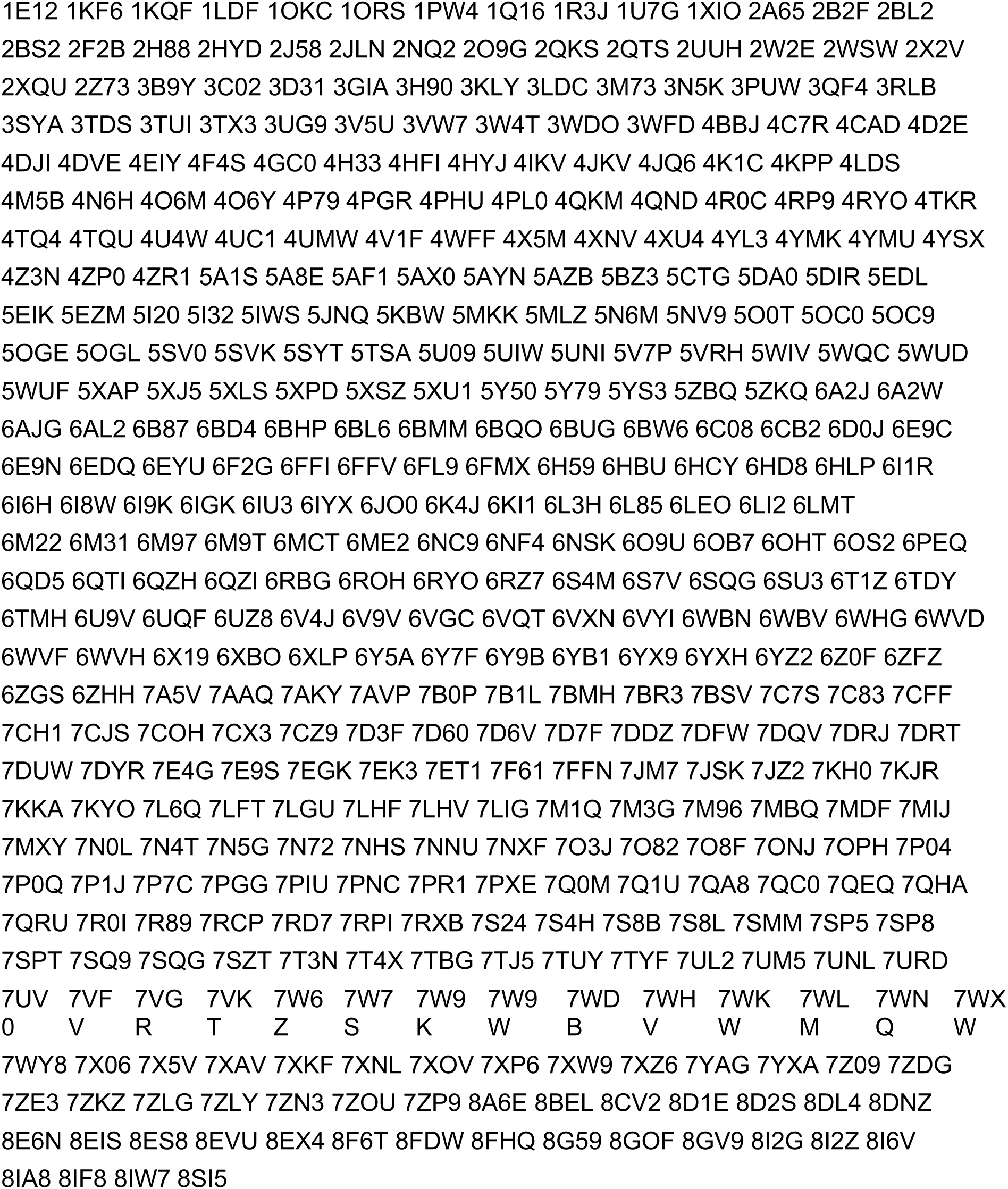
Non-redundant database of membrane protein structures. The transmembrane spans of these membrane proteins were extracted using OPM definitions, and structures were reduced by their internal symmetry into their unique substructures as the reduced database of unique TM tertiary and quaternary structures.

**Supplementary Table 2.**
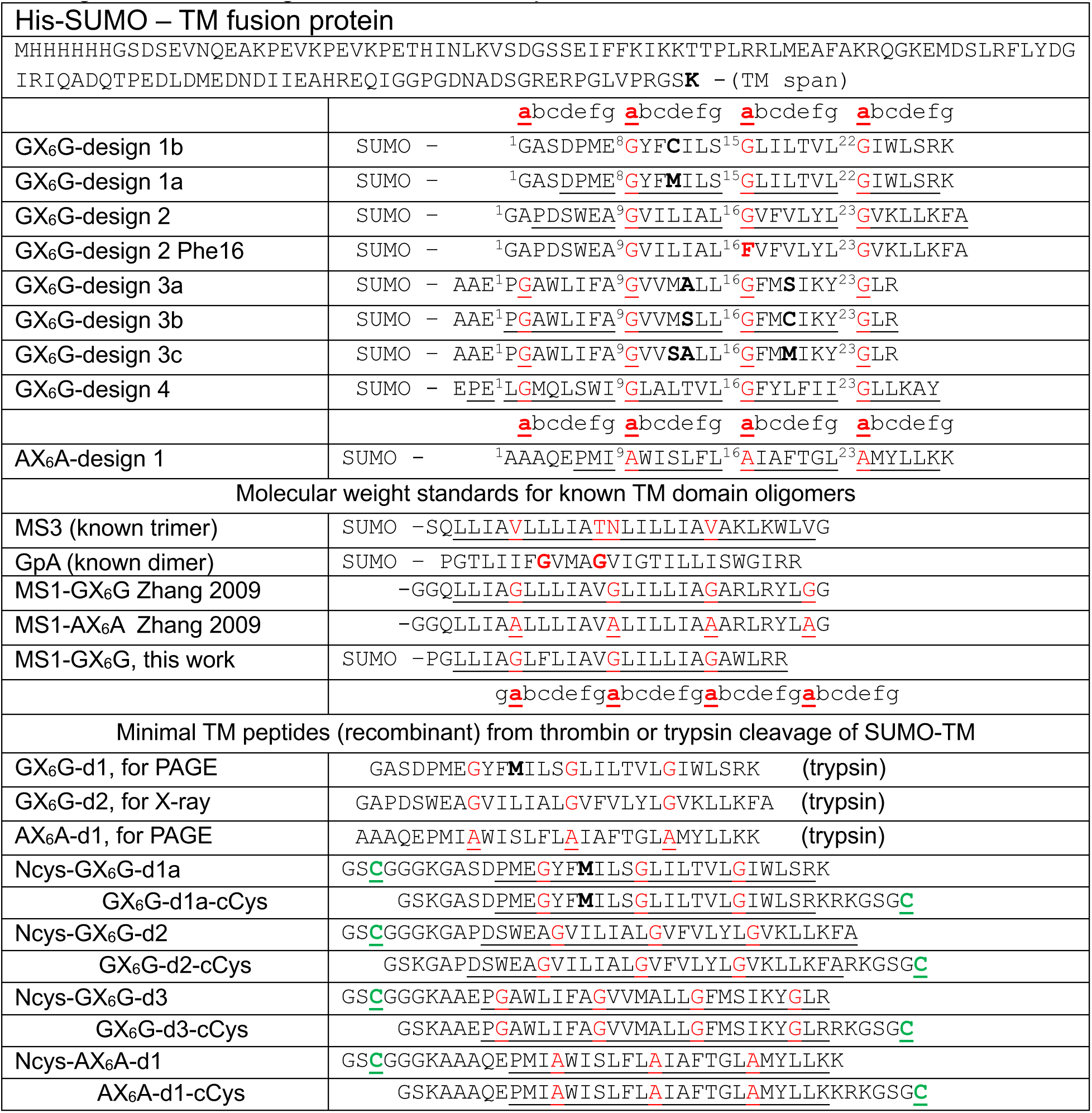
Relevant 1-span protein and peptide sequences in this work. . Designed TM domain sequences as 6xHis-SUMO fusion protein (numbered from 1 after the bolded linker Lys) and TM peptides yielded from cleavage with thrombin (with cysteine, green) or trypsin digestion. Mutation sites bolded. TM regions underlined, aligned by heptad (*abcdefg*). Intended motif small residues, red. TM peptides numbered starting after SUMO region near the first expected transmembrane helix residue

**Supplementary Table 3.** Antiparallel helix-helix geometry of TM designs by coiled-coil approximations. Best fit parameters of the generalized versions of Crick’s coiled-coiled equations from ref ^3^ for crystal structure of GX6G Design-2 or design models of GX6G designs 1, 3, 4, and AX6A Design (final frame of 200 ns MD simulation). Geometry data for the centroid example of the TM building block cluster 6 from ref ^4^ is listed as well.

|  | GX6G Design-1 | GX6G Design-2 | GX6G Design-3 | GX6G Design-4 | AX6A Design | Cluster 6 Centroid |
| --- | --- | --- | --- | --- | --- | --- |
| R0 (Å) | 3.85 | 3.72 | 3.71 | 3.55 | 4.13 | 4.13 |
| R1 (Å) | 2.32 | 2.24 | 2.30 | 2.32 | 2.32 | 2.23 |
| $\omega_0$ (°/res) | -2.61 | -3.22 | -2.87 | -3.4 | -3.74 | -3.61 |
| $\omega_1$ (°/res) | 101.6 | 102.5 | 102.3 | 102.4 | 103.0 | 102.9 |
| $\alpha$ (°) | -6.8 | -8.0 | -7.2 | -8.2 | -10.5 | -10.0 |
| $\phi_1$ (°) | -115.1 | -131.0 | 76.1 | 119.0 | -21.6 | 130.4 |
| $\Delta Z_{off}$ (Å) | 2.1 | -2.6 | 2.5 | -1.8 | 2.0 | 2.4 |
| RMSD CC fit | 0.39 Å | 0.45 Å | 0.35 Å | 0.67 Å | 0.30 Å | 0.61 Å |
| Distance | 7.70 | 7.44 | 7.42 | 7.10 | 8.24 | 8.25 |
| Crossing (°) | -154.6 | -167.6 | -170.3 | -167.7 | -165.3 | -158.11 |

**Supplementary Table 4.** Data collection and refinement statistics for G-X6-G Design-2.

|  | G-X6-G *de novo* Design-2 (pdb: 8SRN) |
| --- | --- |
| **Data collection** |  |
| Space group | C 2 2 2<sub>1</sub> |
| Cell dimensions |  |
| *a*, *b*, *c* (Å) | 54.0, 71.4, 101.1 |
| $\alpha$, $\beta$, $\gamma$ (°) | 90.0, 90.0, 90.0 |
| Resolution (Å) | 35.7 - 3.30 (3.42 - 3.30) |
| *R*<sub>sym</sub> or *R*<sub>merge</sub> | 0.09 (0.498) |
| *I* / σ*I* | 27.5 (2.4) |
| Completeness (%) | 93.82 (78.77) |
| Redundancy | 9.6 (7.4) |
| **Refinement** |  |
| Resolution (Å) | 3.3 |
| No. reflections | 2953 |
| *R*<sub>work</sub> / *R*<sub>free</sub> | 0.253 / 0.254 |
| No. atoms |  |
| Protein | 1034 |
| Ligand/ion | 0 |
| Water | 0 |
| *B*-factors |  |
| Protein | 90.0 |
| Ligand/ion | N/A |
| Water | N/A |
| R.m.s. deviations |  |
| Bond lengths (Å) | 0.005 |
| Bond angles (°) | 0.96 |

\*Data from 1 crystal

**Supplementary Table 5.**
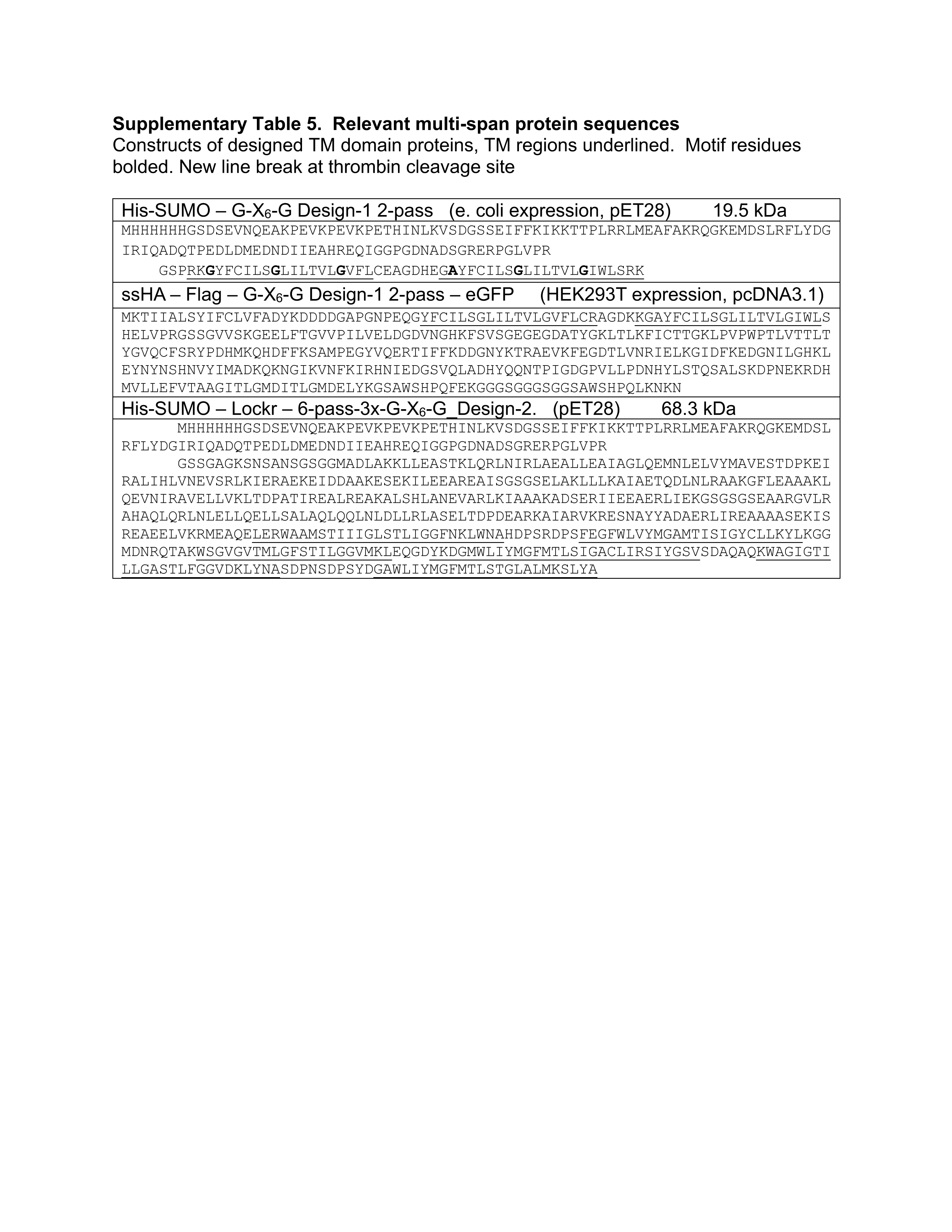
Relevant multi-span protein sequences.

